# Robustness of DNA Looping Across Multiple Divisions in Individual Bacteria

**DOI:** 10.1101/2021.12.28.474367

**Authors:** Chang Chang, Mayra Garcia-Alcala, Leonor Saiz, Jose M.G. Vilar, Philippe Cluzel

## Abstract

DNA looping has emerged as a central paradigm of transcriptional regulation as it is shared across many living systems. One core property of DNA looping-based regulation is its ability to greatly enhance repression or activation of genes with only a few copies of transcriptional regulators. However, this property based on small number of proteins raises the question of the robustness of such a mechanism with respect to the large intracellular perturbations taking place during growth and division of the cell. Here we address the issue of sensitivity to variations of intracellular parameters of gene regulation by DNA looping. We use the *lac* system as a prototype to experimentally identify the key features of the robustness of DNA looping in growing *E. coli* cells. Surprisingly, we observe time intervals of tight repression spanning across division events, which can sometimes exceed ten generations. Remarkably, the distribution of such long time intervals exhibits memoryless statistics that is mostly insensitive to repressor concentration, cell division events, and the number of distinct loops accessible to the system. By contrast, gene regulation becomes highly sensitive to these perturbations when DNA looping is absent. Using stochastic simulations, we propose that the robustness to division events of memoryless distributions emerges from the competition between fast, multiple re-binding events of repressors and slow initiation rate of the RNA-polymerase. We argue that fast re-binding events are a direct consequence of DNA looping that ensures robust gene repression across a range of intracellular perturbations.

**Significance statement:** It is well-established that certain intracellular regulators can stabilize DNA loops to greatly enhance activation or repression of gene transcription. In vitro but also in vivo ensemble measurements have determined that only a few copies of regulators are in fact needed to stably form DNA loops. In view of such a small number, we address the issue of sensitivity of gene regulation by DNA looping to variations of intracellular parameters in individual growing *E. coli* bacteria. Surprisingly, we find that DNA looping from the *lac* system is robust to a range of perturbations including divisions during which cells can maintain tight repression over many generations. We propose a mechanism that governs the observed robustness across a range of intracellular perturbations.

## Introduction

Several genetic systems in bacteria are known to use only few repressors to maintain low levels of expression, such as the *lac*, arabinose and lysogenic regulation (1-3). The diversity of these systems underlines the importance for the cell to have selected certain molecular mechanisms for efficiently maintaining low expression levels together with low level of repressors. The *lac* operon is arguably among the most studied genetic regulatory systems of this class and is known to utilize a higher-order structure of DNA, DNA looping, to repress efficiently the activity of the *lac* promoter using only a handful of copies of repressors (1, 4-6). While the strong repression mediated by DNA looping have clearly been established in vivo and in vitro, the fact that it relies on a small number of repressors to function, however, makes this molecular mechanism potentially sensitive to intracellular perturbations. For example, a small number of repressors can fluctuate greatly at cell division, which may yield undesirable promoter leaks (7-9), and it is still an open problem to know whether DNA looping can maintain repression even across several divisions. Indeed, a standard assumption is that gene duplication and cell division may disrupt the looping structure and binding of the repressors to DNA (10), which would consequently limit the duration of repression intervals. Moreover, during cellular growth, DNA replicates and gene dosage increases as a function of time, which may dynamically change the ratio of the number DNA binding sites with that of repressors.

In light of these outstanding questions, we aimed at quantitatively characterizing how robust the repression of DNA looping is with respect to intracellular perturbations in individual growing bacteria. In our experiments, we monitor the spontaneous leakiness of the promoter, as a measure for the repression level of the *lac* promoter in the presence or absence of DNA looping. Using a microfluidic device, we record long time series associated with the leakiness of the *lac* promoter in individual growing *E. coli* cells across more than 40 generations. We use as a starting point the model by Vilar and colleagues that proposed that the change of free energy associated with DNA looping formation is equivalent to the existence of a very large ‘local’ repressor concentration, effectively hundred times larger than the ‘global’, wild type repressor concentration (11, 12). One key prediction of this model is that repression by means of DNA looping is robust to fluctuations of repressors concentration, while repression in the absence of DNA looping is sensitive to fluctuations. We combine several techniques to record and analyze the spontaneous leakiness of the *lac* system (Fig. 1).

**Fig. 1.**
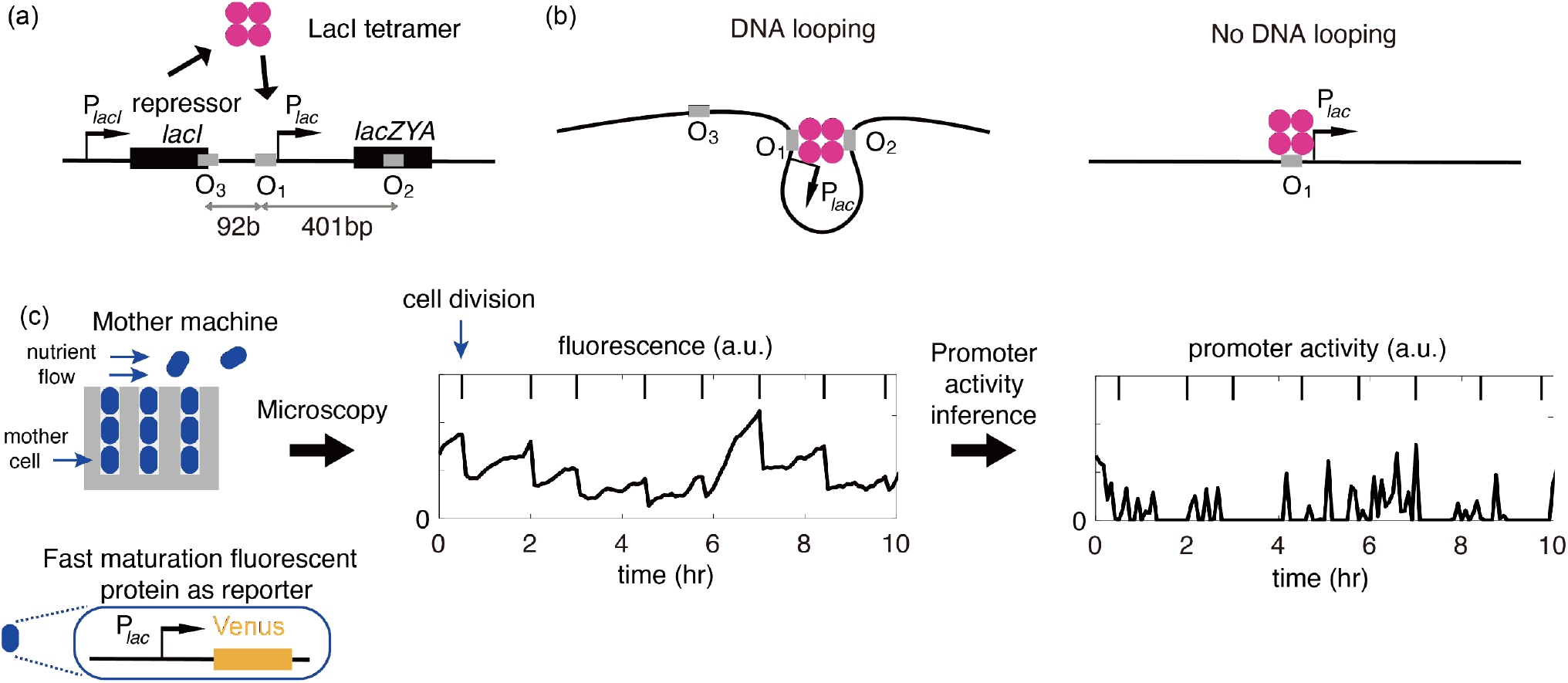
Monitoring multi-generational leakiness of the endogenous *lac* promoter in single cells. (a-b) Wild type *lac* operon maintains a low level of expression by the means of DNA looping. (a) Genetic organization of the *lac* system, where O_i_ denote the operators *i*, and are DNA specific sequences where the repressor LacI binds. LacI is fully functional as a tetramer (19), and can bind to any of three possible operators. (b) Cartoon that illustrates how several operators can mediate the formation of DNA looping while one operator alone cannot. (c) We developed an experimental platform to monitor promoter activity for low expression systems across multiple cell divisions.

## Experimental design

Due to the very low leaking rate, we monitor the promoter activity from single cells and across many division cycles using a microfluidics device, called the ‘mother machine’ (13, 14). In this device, cells grow under chemostatic conditions. The mother cell is trapped at the bottom of a microfluidic channel, while daughter cells are washed away when they exit the channel. In our experiments, to estimate promoter activity we use the production rate of a fluorescent reporter driven by a copy of the *lac* promoter. To measure gene expressions with an improved temporal resolution and signal-to-noise ratio, we use a fast maturating fluorescent protein (VenusNB, maturation half-time 4.1 ± 0.3 *min* (15)), together with an optimized ribosome binding site to maximize the yield of translation of the fluorescent reporter (16). Maximizing the yield of translation helps us to detect small transcriptional bursts that are usually not directly detectable at the single cell level. It was found earlier that a large fraction of cells do not contain even one copy of the fluorescent protein controlled by the *lac* operon (17), thus the fluorescent levels of most of the cells is often nearing the level of auto-fluorescence. Under this condition of such a low expression level, we further developed a probabilistic algorithm that explicitly takes into account the fluctuations of auto-fluorescence background to robustly discriminate promoter activity from background noise (Fig. S2 and SI).

Cells were cultured overnight in M9 medium with 0.4% glycerol as carbon source, then loaded and cultured in exponential growth by steadily flushing them with fresh media in the mother machine (≥ 40 hours; 30°C; also see SI). Phase contrast and fluorescence images (Zeiss Axiovert 200M microscopy) were captured for each field of view with a dwell time of 5 minutes (15). We use an open-source software Molyso (18) to perform cell segmentation and lineage tracking, and further customized the codes of that software for proofreading (see SI).

### Statistics of promoter leakiness with and without DNA looping

The regulatory regions of the *lac* operon consist of one main operator O_1_, and two auxiliary operators O_2_ and O_3_ (Fig. 1a). When a tetramer of LacI repressor simultaneously bind two operators, e.g. O_1_-O_2_ or O_1_-O_3_, it can form a stable DNA loop (Fig. 1b, left panel). Previous population measurements suggest that with only ∼10 repressors, the *lac* operon can maintain a repression level ∼100 times stronger than in the absence of DNA looping when there is only the single operator, O_1_, present (1, 20). To compare the statistics of leaky events with and without DNA looping, we investigated two *E. coli* strains: one that carries all three operators, denoted as the *Loops* strain; while the other one only carries the main operator O_1_, denoted as the *No-loop* strain (Fig. 1b and Table S1). Without DNA looping, the promoter exhibits frequent transcriptional bursts (Fig. 2a, left panel). By contrast, in the presence of DNA looping, the promoter leaks unfrequently, and transcriptional bursts are separated by very long periods of time that can exceed sometimes ten cell cycles (Fig. 2a, right panel).

**Fig. 2.**
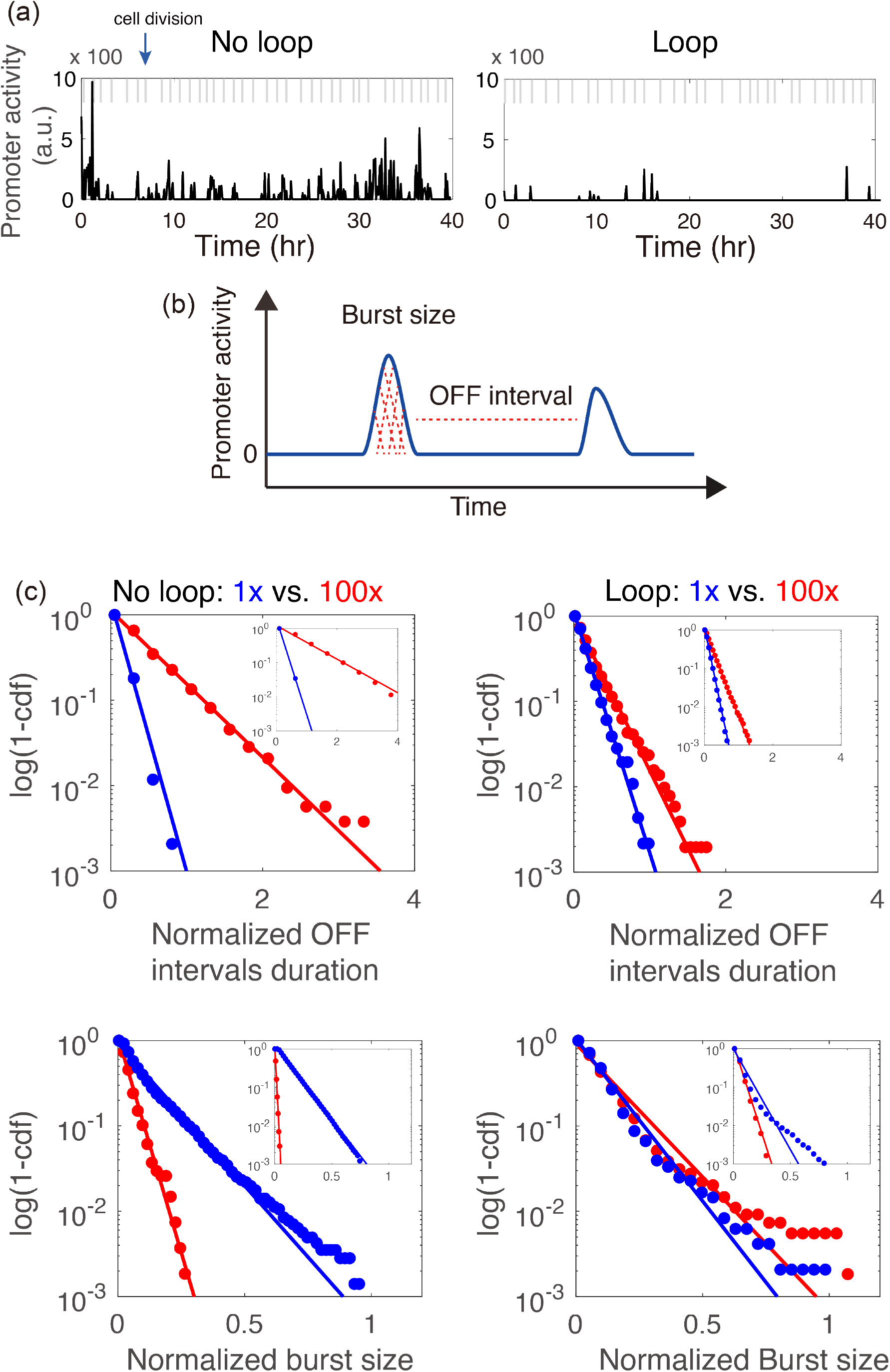
Statistics of promoter activity with and without DNA looping in single cells. (a) Example of promoter activity from single-cell traces of (left) the No-loop and (right) the Loops strain across multiple cell divisions. (b) We use two key physical quantities to characterize the dynamics of promoter activity: the duration of OFF intervals and the burst size of pulses. (c) Cumulative distributions (*P*(*X* ≥ *x*)) of statistics with various repressor concentrations (denoted by color). Dots represent the statistics from experiments and lines give the fitting (linear fits in semi-log space), and insets give the statistics and the fitting from simulated time series in the absence of cell division using Vilar *et al*. model (see Fig. S14 for more details). In each panel and each inset, the statistics are normalized with the maximum value from the blue dots so that the blue curve ends around 1. Normalization factors: Loops strains (OFF interval=1440 min, burst size=2248 a.u.), No-loop strains (OFF interval=395 min, burst size=5359 a.u.). Slopes of the fitting of OFF intervals before normalization are given as follows: Loops (−0.00446 *min*^−1^), 100x/Loops (−0.00289 *min*^−1^), No-loop (−0.01838 *min*^−1^), 100x/No-loop (−0.00497 *min*^−1^).

We characterized the dynamics of the promoter activity using two quantities: the duration of OFF intervals and the transcriptional bursts size (Fig. 2b; # lineages ≥ 50). Surprisingly, we find that the OFF intervals from the Loops strain follows an exponential distribution like for the simpler No-loop strains (Fig. 2c). A memoryless statistical process was not expected for the Loops strain, because the OFF intervals were on average longer than several cell cycles, and complex statistics would have been more in line with the multiple steps processes that accompany cell division. On the other hand, the burst size of the Loops and No-loop strains have a linear region in the semi-log space but followed by a long tail (Fig. 2c). The Loops strain overall exhibits significantly longer OFF intervals (OFF mean = 202 min, standard error [SE] ±10, or on average 2.8 cell cycles) and smaller burst size (213 [SE] ±11 a.u.) compared to the No-loop strain (OFF = 47± [SE] 1 min or 0.6 cell cycles, burst size = 592 [SE] ±18 a.u.).

We further investigate how robustly DNA looping could repress the promoter versus using a strain that has only the main operator O_1_, both in the presence of high concentration of repressors. We performed this experiment to test the key prediction of Vilar *et al*. model, i.e., the promoter leakiness is insensitive to repressor concentration in the presence of DNA looping. Consequently, we constructed additional two strains, *100x/Loops* and *100x/No-loop*, where the concentration of repressors is ∼100 times larger than that in the Loops and No-loop strains (SI). Under those conditions, the OFF intervals still follow exponential distributions (Fig. 2c). Again, the burst size of 100x/Loops has a long tail, but not the 100x/No-loop strain. However, in the 100x/No-loop strain promoter leakiness is very sensitive to the increase of the repressor, with OFF intervals increasing to 178 [SE] ±8 min (or 2.6 cell cycles) and burst size to 210 [SE] ±9 a.u.. By contrast, 100x/Loops is insensitive to the increase in LacI repressor concentration and show only slightly longer OFF intervals (292 [SE] ±15 min or 4.5 cell cycles) than that of the Loops strain. As for the distributions of the burst size, they are similar (223 [SE] ±13 a.u.), indicating the typical burst size of the Loops strain has already been reduced to its lowest limit, which we reasoned may be associated with the synthesis of only one mRNA per pulse. The burst size of the 100x/No-loop strain also reached a similar limit in the presence of high repressor concentration. Our first observations are in agreement with the predictions of the Vilar *et al*. model.

That OFF intervals across many divisions in the Loops strains follow exponential distributions is unexpected, because it is the signature of a memoryless one-step process associated with a single rate. We hypothesize that this one-step process that control the statistics of OFF intervals in the Loops strains is largely dominated by the unbinding of repressors from the operator O_1_, regardless of whether the auxiliary operators O_2_ and O_3_ are bound or not.

To check this hypothesis, we knocked out one additional operator (O_2_ or O_3_) in the Loops strains and found that the OFF intervals of those strains that can form only one loop (either O_1_-O_2_ or O_1_-O_3_) follow exponential distributions as well (Fig. S13). Furthermore, we observe that the O_1_-O_2_ One-loop strain exhibit qualitatively similar timescales of the OFF intervals (204 [SE] ± 9 min) as for the Loops strain that have the possibility to form multiple alternative loops (Fig. S13). This result indicates that the switching between different loops is rare and hardly modifies the statistics for OFF intervals.

To evaluate the impact of cell division on the OFF intervals, we performed stochastic simulations of gene expression using the Vilar *et al*. model (12) (also see SI). Modeling cell division by periodically forcing the unbinding of the repressor from the operators is unable to reproduce the long and exponentially distributed intervals, as well as the insensitivity to repressor concentration in the Loops strain (Fig. S14). By contrast, stochastic simulations in the absence of cell division can reproduce the experimentally observed statistics (Fig. 2c, insets), indicating that DNA looping is robust to the perturbations associated with cell division.

### A theoretical model without cell divisions

Most observations in our experiments can be qualitatively understood with a three-state model extended from Vilar *et al*. model (11, 12): (I) State *B*: the repressor LacI is bound to O_1_; (II) State *E*: the operator O_1_ is empty and freed from RNA polymerase; and (III) State *TS*: O_1_ is cleared from RNA polymerase and transcription starts (regardless of the states of O_2_ or O_3_). We further assume that only one transcript is produced in state *TS* and that the system returns to state *E* immediately. The transitions between states are described by

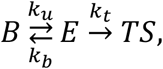

where *k*_*b*_ is the effective binding rate for the repressors to O_1_, *k*_*u*_ gives the unbinding rate for a repressor from O_1_ (0.10 min^−1^), and *k*_*t*_ is the effective transcription rate (20 VenusNB ⋅ min^−1^). Without DNA looping, repressors follow a simple ON-OFF dynamics, thus *k*_*b*_ scales linearly with the repressor concentration *n*_*R*_ (*n*_*R*_ = 10 molecules per cell in 1x LacI strain and *n*_*R*_ = 1000 in 100x LacI). For a strain with DNA looping, when the system is in state *E*, most likely one of its auxiliary operators is with a repressor, given the free energy difference between a bound and a free operator. For a *E*-*B* transition, either a repressor from the rest of the cell, denoted as ‘global’, binds to O_1_; or the repressor that has already bound to an auxiliary operator binds rapidly to O_1_, denoted as ‘local’. The free energy difference of the looping formation, 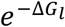, is effectively equivalent to a very large ‘local’ concentration *n*_*L*_ (*n*_*L*_ = 0 for No-loop, 1080 for Loops) (11, 12). Considering both situations, we have *k*_*b*_ = *k*_*on*_ (*n*_*R*_ + *n*_*L*_) where *k*_*on*_ is the binding rate for a single repressor (0.28 moiecuie^−1^min^−1^).

An OFF interval consists of one or multiple rounds of *E*-*B* transitions before the system goes to state *TS*. The probability of *l* rounds to occur is *P*_*l*_= *α*^*l*−1^(1 − *α*), with 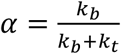 be the probability of entering state *B* from *E*. Considering that *k*_*u*_ ≪ *k*_*b*_, as implied by the physical parameters of the system, the timing at which *l* unbinding events happen is given by the composition of *l* exponential decays, which can be described by the Erlang distribution, 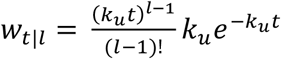, resulting in a distribution of waiting times between transcriptional events (OFF intervals) 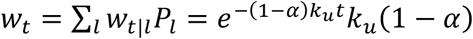. Thus, the average duration of the OFF interval is 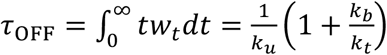. Theoretical calculations suggests that the ratio of *τ*_*off*_ between the Loops strains with 1x and 100x repressor concentration is ∼2, but that ratio between the No-loop strains is ∼14 (see SI). Consequently, the model qualitatively predicts the great sensitivity of the No-loop strains and the insensitivity of the Loops strains to repressor concentration.

On the other hand, the burst size of a pulse is proportional to the number of *E*-*TS* transitions before the system goes to state *B*, equivalently the number of transcripts. Starting from state *E*, the probability of entering state *TS* is 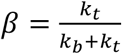. The probability to produce *r* transcripts in a pulse is *P*_r_ = *β*^*r*−1^(1 − *β*), a geometric distribution with an average number 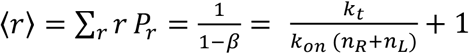. When *n*_*R*_ + *n*_*L*_ is large, ⟨*r*⟩ → 1. Theoretical calculations suggest the No-loop strain is expected to have more than one transcript per pulse (estimated as ∼8), by contrast, the other three conditions with DNA looping or high concentration of repressors are predicted to have only about one transcript per burst. These predictions are in line with our experimental observations showing that the bursts size cannot be reduced further in the Loops strain even when we drastically increase the repressor concentration.

### Correlation between promoter activity and gene dosage during growth

As described by the Cooper-Helmstetter relation (21), gene expression depends on global factors such as gene dosage (22). As the cell grows, DNA replicates in such a way that the average copies of chromosome is maintained after division, but between two division events the gene dosage increases. In *E. coli*, it has been reported that the promoter activity of a gene with a high expression level is correlated with the phase of cell cycle (23) and has been quantitatively measured in Ref. (24). Under induction (the removal of the repressors), the gene expression is constitutive, and the promoter activity of the *lac* operon exhibits a flat region at the early phase of cell cycle, and gradually increase to about twice its initial level (24).

Next, we monitored how the spontaneous leakiness of the repressed promoter correlates with the cell cycle in the presence and absence of DNA loops. Although the absolute promoter activity greatly varies across our four strains, e.g. the promoter activity of the No-loop strain is ∼10 times larger than that of the Loops strain, they all show positive correlations with cell cycle progression (Fig. 3, inset). Furthermore, after normalizing by the mean promoter activities, all curves with different expression level collapse (Fig. 3), indicating that the promoter activity in all strains have the same dependence on the cell cycle in the presence or the absence of DNA looping. We interpret the increase of promoter activity within the cell cycle as a consequence of an increase of gene dosage due to DNA replication that is similar across all our strains.

**Fig. 3.**
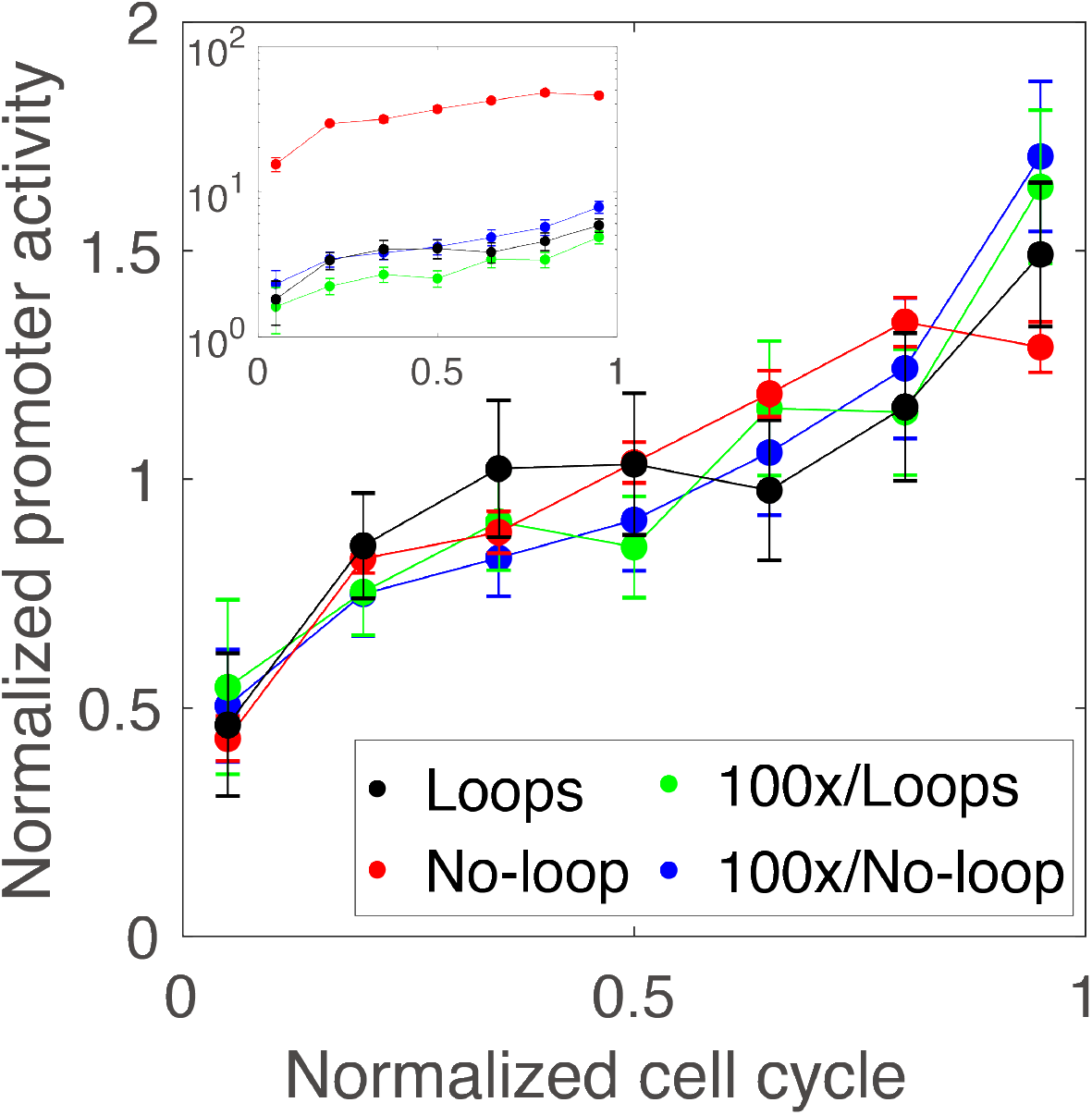
Promoter activity as a function of time span between two cell-division events. Time span between two successive division events is normalized by the total duration of this interval. The normalized averaged promoter activity is obtained by dividing the promoter activity of each strain by its mean. Error bars represent the standard error. The inset gives unnormalized promoter activity for each strain.

## Discussion

The extension of Vilar *et al*. model and stochastic simulation without considering DNA replication and cell division is able to reproduce several key observations from the experiments, including memoryless distributions of the OFF intervals and extremely long OFF intervals with DNA looping, as well as its insensitivity to repressor concentration. Given the repressor LacI will be removed during DNA replication (10), the DNA looping structure may be affected and the multi-generational suppression of promoter activity is unexpected, especially in present of very few repressors (∼ 10; see Ref. (25)). Meanwhile, in a cell cycle, the gene dosage of both repressor and *lac* operon doubles. During a cell division, the number of the *lac* operon is exactly halved, but the number of the repressors that a daughter cell inherits fluctuates. Remarkably, neither the variations due gene dosage within cell cycles nor perturbations associated with division events interrupt the long OFF intervals and their associated simple exponential distribution.

Using the assumption of ‘local’ repressor concentration from Vilar *et al*. model, we reason that the rebinding of the repressor onto the operator after gene duplication is fast. The *ab initio* search time for LacI to bind free O_1_ is >30 seconds (26, 27), but it only takes ∼3 seconds for a polymerase to start transcription (28). In the presence of DNA looping, if rebinding of the repressor were slower than the polymerase initiation rate, the duration of repression intervals would be of the order of the division time, like in the No-loop strain. Thus, the removal of repressors from O_1_ during DNA replication adds only a few more rebinding ‘events’ per cell cycle. Consequently, in presence of DNA looping, division events should not affect the memoryless statistics of the OFF intervals.

The time scale for long repression intervals observed in our experiments can be reconciled with the short lifetime of DNA loop measured in vitro (29). Using Chen *et al*. measurements (29) and Refs. (30, 31), we estimated that the in vivo loop lifetime was of the order of 10^3^ sec. and that of the open loop was ∼1 sec.. These values are similar to those used in our theoretical model that provides a simple mechanism for the existence of OFF intervals longer than division cycles. When O_1_ is unoccupied, we hypothesized that there is competition between RNA polymerase initiation and repressor rebinding events. However, the typical timescale for the RNA polymerase initiation (∼3 sec.) is about an order of magnitude slower than the timescale to re-form a loop (∼0.1 sec., estimation from our model), consequently transcription initiation happens only once every 30 attempts.

Additionally, we use the Cooper-Helmstetter relation based on growth rate and find under our condition that the number of the lac operon copies averaged over a division cycle is about two. Therefore, the number of replication events that potentially perturb repressor binding is very small, and we reason that DNA replication and division should not significantly affect the mean duration of the OFF intervals. Indeed, we shall expect the mean OFF intervals in the presence of DNA looping to be in the order of 30 times of the loop lifetime (10^3^ sec) measured in vitro, i.e. about 7 cell cycles in line with our experiments.

Cai *et al*. (2006) measured the burst frequency of the *lac* operon (0.11 ± 0.03) using a *β*-gal assay combined with a microfluidic device, averaged over multiple cells without across cell cycles (17). They also inferred this burst frequency (0.16 ± 0.05) from a population distribution of *β*-gal, based on a steady-state theory derived from a master equation (17). Our direct measurement of a cell with DNA looping over multiple cell cycles (0.36±0.11) is larger than both estimations.

In summary, we report the robustness of DNA looping to intracellular perturbations across multiple cell cycles. While a small copy number of repressors are present in the cell, we find repression with DNA looping insensitive to the variations of intracellular environment, such as repressors concentration, cell divisions, and detailed configurations of DNA loops. We speculate that similar robustness plays a crucial rule in other genetic regulatory systems beyond the *lac* operon.

## Author Contributions

C.C. and P.C. designed research; C.C. performed experimental research with the help from M.G.A.; C.C. and P.C. analyzed the data, performed statistical analyses, and designed the inference algorithm; L.S. and J.M.G.V. designed and developed theoretical approaches and performed stochastic simulations. C.C., L.S., J.M.G.V. and P.C. wrote the paper with the editing from M.G.A..

## Acknowledgments

We thank Guangwei Si, Junhao Zhu, Richard Losick, J. Mark Kim, Kritika Gupta, and Jimmy Martenson for fruitful discussions. The authors are indebted to Enrique Balleza who performed preliminary experiments for this study. This work was supported by NSF Grant 1615487 to P.C.. J.M.G.V. acknowledges support from Ministerio de Ciencia e Innovación under grant no. PGC2018-101282-B-I00 (MCI/AEI/FEDER, UE). L.S. is indebted to the MIT Institute for Medical Engineering & Science for hosting her for a sabbatical stay during the early stages of this work.

