## Supplementary Information for "Robustness of DNA Looping Across Multiple Divisions in Individual Bacteria"

**Genetic cloning**
All clones were derived from *Escherichia coli* strain MG1655 (The Coli Genetic Stock Center, Yale University; CGSC 6300). The carbon source in our experiments is glycerol, under which condition the synthesis of flagella is induced. As a consequence, many of the cells in the mother machine would swim away from the channels of the mother. To disable the motility of the cells, we knocked out in all strains of this study *fliC*, the gene that encodes for the protein flagellin so that flagellar filament cannot be synthetized.

We constructed all strains using the lambda red recombination technique (1, 2), with the helper plasmid pSIM5 expressing recombineering functions (3). To construct a strain without an antibiotic marker, we used a "scarless" chromosomal engineering technique based on a counter-selection cassette (4). In the first step, we inserted a linear fragment from $P_{araB}$-*ccdB* cassette (gift from J. Mark Kim) into target site using kanamycin for selection. Then this region was further replaced by the final construct, using arabinose for selection. The expression of CcdB is toxic to *E. coli* in the absence of CcdA, thus, with the induction by arabinose, the fitness of a cell without $P_{araB}$-*ccdB* cassette (e.g., replaced by the final construct) outperforms a cell with the cassette. On the other hand, to construct a strain with an antibiotic marker, we directly integrated into the target site a linear fragment with both the final construct and the antibiotic marker, using this specific antibiotic for selection (kanamycin in our study). Most of the linear fragments were derived from plasmids (Table S2). For all strains used in mother machine experiments, the temperature-sensitive helper plasmid pSIM5 was removed (3), and their single clones were obtained for further experiments. The code and the descriptions of the strains are available in Table S1.

We constructed CC-41 from MG1655 by introducing Δ*fliC* background in a "scarless" way (Table S1), which is called the background strain that does not carry any fluorescence reporter gene. The background strain is the starting point of all other derived strains.

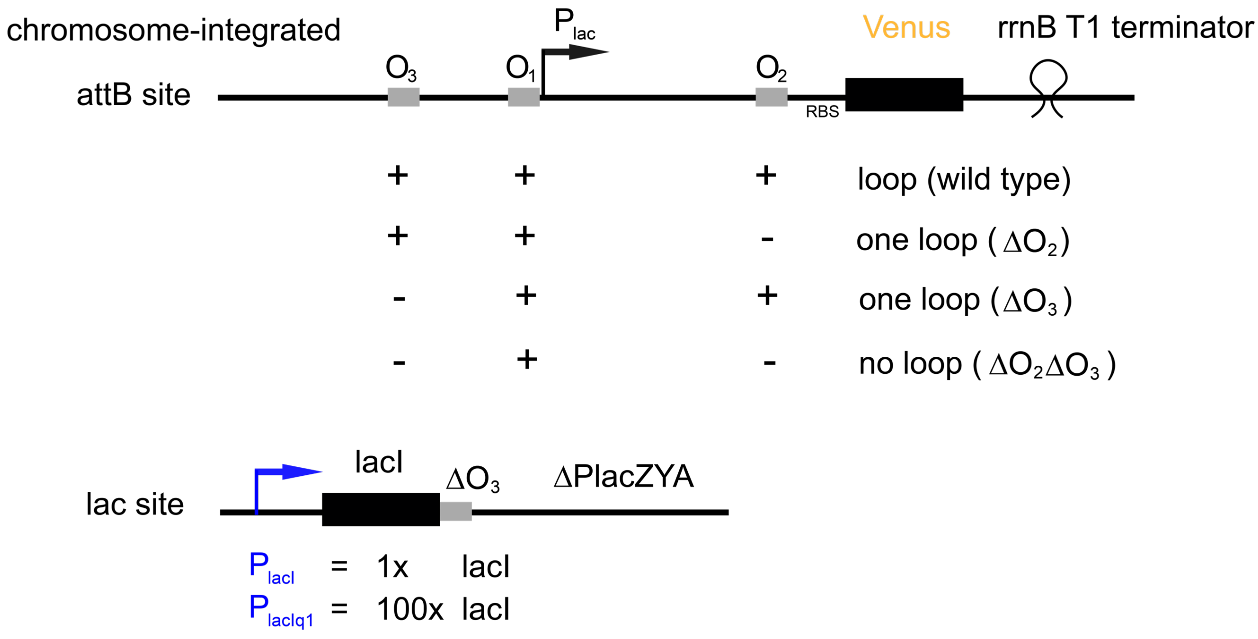

**Figure S1. Cloning design.** All constructs are integrated into the chromosome rather than a plasmid to avoid cell-to-cell copy number variations. The native *lacI* site is decoupled from the DNA looping cassette site that was cloned into the *attB* site. At the *lac* site, $O_3$ partially overlaps with the 3' end of *lacI* that was kept intact after the $O_3$ deletion (Table S3), and no antibiotic marker is integrated. At the *attB* site, *lacZ* is truncated at ~ 50 base pair after $O_2$, following by the fast maturating fluorescence reporter VenusNB (5) that is under the control of $P_{lac}$ and is followed by a kanamycin resistant gene (after the *rrnB* T1 terminator).

In this study, we want to explore the effect of the number of functional operators in the DNA looping cassette and of the repressor concentration on repression strength of the *lac* operon. To decouple those two factors, we kept the *lacI gene* intact at the native site but deleted the looping cassette from the native site and cloned it at the *attB* site (Fig. S1). First, we constructed CC-50 from CC-41 using a linear fragment from $P_{araB}$-*ccdB* cassette to knock out all genes between *lacI* and *lacA* at the *lac* site. Secondly, we inserted *lacI* with different promoters back to the *lac* site ($P_{lacI}$ for CC-51 that has wild type repressor expression, and $P_{lacIq1}$ for CC-53 that over-expresses LacI at an order of 100 times (6)) but without DNA looping cassette nor no antibiotic marker. In a final step, CC-54, CC-58, CC-64 and CC-67 were constructed from CC-51 and CC-53 by cloning the DNA looping cassette to the *attB* site, and those strains consist of high and low LacI repressor expression combined with either the presence or absence of the DNA looping cassette (the DNA looping cassette can be disabled by knocking out *lacO*$_2$ and *lacO*$_3$; see Table S3). For simplicity, we denote CC-54 as the *Loops* strain, CC-58 as the *100x/Loops* strain, CC-64 as the *No-loop* strain, and CC-67 as the *100x/No-loop* strain (Table S1), where the number before the slash refers to the order of the relative LacI concentration to the wild type. In addition to those Loops and No-loop strains, we constructed four One-loop strains ($O_1$-$O_2$ One-loop, $O_1$-$O_3$ One-loop, 100x/$O_1$-$O_2$ One-loop, 100x/$O_1$-$O_3$ One-loop; see Table S1 and Fig. S13).

| Strain | Parent strain | Genotype | Notation used in the text |
|---|---|---|---|
| CC-41 | | *E. coli, str.* MG1655, Δ*fliC* | background |
| CC-50 | CC-41 | (Δ*lacI-lacA*)::$P_{araB}$-*ccdB* | |
| CC-51 | CC-50 | Δ(*lacO*$_3$-*lacA*) | |
| CC-53 | CC-50 | Δ$P_{lacI}$::$P_{lacIq1}$, Δ(*lacO*$_3$-*lacA*) | |
| CC-54 | CC-51 | *attB*::$P_{lac}$-*lacZ*(1-430)-SD-*mVenusNB* | Loops |
| CC-58 | CC-53 | *attB*::$P_{lac}$-*lacZ*(1-430)-SD-*mVenusNB* | 100x/Loops |
| CC-62 | CC-51 | *attB*::$P_{lac}$-*lacZ*(1-430)-Δ(*lacO*$_2$)-SD-*mVenusNB* | $O_1$-$O_3$ One-loop |
| CC-63 | CC-51 | *attB*::$P_{lac}$-Δ(*lacO*$_3$)-*lacZ*(1-430)-SD-*mVenusNB* | $O_1$-$O_2$ One-loop |
| CC-64 | CC-51 | *attB*::$P_{lac}$-Δ(*lacO*$_3$)-*lacZ*(1-430)-Δ(*lacO*$_2$)-SD-*mVenusNB* | No-loop |
| CC-65 | CC-53 | *attB*::$P_{lac}$-*lacZ*(1-430)-Δ(*lacO*$_2$)-SD-*mVenusNB* | 100x/$O_1$-$O_3$ One-loop |
| CC-66 | CC-53 | *attB*::$P_{lac}$-Δ(*lacO*$_3$)-*lacZ*(1-430)-SD-*mVenusNB* | 100x/$O_1$-$O_2$ One-loop |
| CC-67 | CC-53 | *attB*::$P_{lac}$-Δ(*lacO*$_3$)-*lacZ*(1-430)-Δ(*lacO*$_2$)-SD-*mVenusNB* | 100x/No-loop |

**Table S1. *E .coli* strains in this study.** The source of all strains is from this work.

| Plasmid | Genotype/Description | Antibiotics | Source |
|---|---|---|---|
| pCC-12 | sc101, attB1-$P_{lac}$-*lacZ*(1-430)-SD-*mVenusNB*-attB2 | Kan | This work |
| pCC-16 | sc101, attB1-$P_{lac}$-Δ(*lacO*$_3$)-*lacZ*(1-430)-Δ(*lacO*$_2$)-SD-*mVenusNB*-attB2 | Kan | This work |

**Table S2. Plasmids in this study.**

| Operator | Sequence |
|---|---|
| lacO$_2$ | GGTTGTTACTCGCTCACATTT |

| | |
|---|---|
| $\Delta lacO_2$ | GGCTGCTATAGCTTGACGTTT |
| $lacO_3$ | GGCAGTGAGCGCAACGCAATT |
| $\Delta lacO_3$ | GGCAGTGATGAAGCTTGTCAG |

**Table S3. Operator and their sequences (7).**

| Site | Primer | Sequence |
|---|---|---|
| *lac* | mCC-1 | AGCAAAACAGATCGAAGAAGGG |
| | mCC-9 | GGTCAAAGAGGCATGATGCGAC |
| *attB* | mCC-46 | AAGACCGCAGAGCAGAGAAC |
| | mCC-47 | TGTTGTCACCTGCTACGACC |
| *fliC* | mCC-98 | GTTGCCGTCAGTCTCAGTTAATCAGGTTAC |
| | mCC-99 | ACCCGACTCCCAGCGATGAAATAC |

**Table S4. Primers used in this study.**

We performed Sanger sequencing to verify these strains (Table S4). We used mCC-1 and mCC-9 for checking the *lac* site, mCC-46 and mCC-47 for the *attB* site, and mCC-98 and mCC-99 for the *fliC* site, verify all news strains with sequencing. Sequencing primers are properly chosen to make sure that all regions related to experiments were examined.

**Experimental setup**

M9 media was prepared with M9 minimal salt (BD, Difco, catalog number: 248510), complemented with 0.1 mM $CaCl_2$ (MilliporeSigma, catalog number: EM1.02378.0500), 2mM $MgSO_4$ (Sigma-Aldrich, catalog number: M1880), 1μg/ml Thiamine (Sigma-Aldrich, catalog number: T4625), 0.85g/L Pluronic F-108 (Sigma-Aldrich, catalog number: 542342), 0.50% casamino acids (BD, Bacto, catalog number: 223050), and 0.40% Glycerol (VWR, BDH, catalog number: BDH1172).

We used Zeiss Axiovert 200M microscope, with all the settings be identical as in (5). In preparation for a typical experiment, cells were grown in M9 media for overnight at 30°C (no antibiotics for the background strain, and 25 ug/ml kanamycin for Loops, One-loop and No-loop strains). At the day of the experiment, the cell culture was centrifuged and loaded into the inlets of microfluidics device by pipetting. Syringes with M9 media (no kanamycin) were connected to the inlets (BD, 60ml), and the outlets were connected to an empty beaker. The initial flow rate was 35 μl/min (~ 1 hour) for cleaning the inlets and outlets and the flow rate was adjusted to 7 μl/min during the experiment. All experiments were performed at 30°C. In this study, one mother machine microfluidics device (8, 9) has four independent quadrants, each has a series of growth channels allowing the observation of the old-pole mother cell and its progeny. This device allows the simultaneous observation of up to four different conditions, each for one quadrant. In each experiment, we included the background strain as a control, and we filled the other quadrants with Loops, One-loop or No-loop strains. Phase contrast and fluorescence images were obtained for each field of view every 5 minutes. After loading the microfluidic device with bacteria, the first five hours of data is discarded to ensure that the cells under observation are in the exponential phase, and subsequent duration of the recorded data is $\geq 40$ hours. The number of lineages of a Loops, One-loop or No-loop strain included in the analysis is $\geq 50$, and the number of lineages of the background strain in the same experiment is $\geq 20$.

**Image processing**

We analyzed our microscopy images based on the software of *molyso*, which includes image registration, cell segmentation and lineage tracking (6). Here, we only tracked the old-pole mother cells, which always stayed at the end of growth channels. We modified the codes of *molyso* to enable manual corrections of the results of cell segmentation. Cell divisions were determined automatically based on cell length, complemented with manual check.

We calculated the total fluorescence per cell as the sum of the fluorescent intensities of all pixels belonging to a cell, subtracted with local background level. We also analyzed a cell without a fluorescent reporter and calculated its total auto-fluorescence.

**The definition of repression**

In our experiments, we monitor the spontaneous leakiness of the promoter, as a measure for the repression level of the *lac* promoter in the presence or absence of DNA looping. Our definition of repression differs from that of Müller-Hill et al. in Refs. (10, 11) that includes the fully induced promoter, however, they both yield the same qualitative pictures.

**Characterizing background fluorescence signals**

With the tight regulation of the DNA looping, most cells contain only few copies of the protein under the control of the *lac* promoter (12). As expected, from our experiments, the distributions of total fluorescence per cell of the Loops and No-loop strains show a low mean in such a way that they are not clearly separated from the distributions of the total auto-fluorescence per cell of the background strain (Fig. S2). This result indicates that, we need to explicitly consider the contribution of the auto-fluorescence to the total measured fluorescence signal in order to identify which part of the measured fluorescent signal is actually directly associated with bursts of VenusNB synthesis and not random fluctuations of auto-fluorescence.

The total auto-fluorescence per cell of the background strain is used for the inference of promoter activity for the Loops and No-loop strains. To determine if the auto-fluorescence signals from different experiments are consistent, we compared the distributions of the auto-fluorescence levels from two repeated experiments. Although existing small differences, they overlap well with each other (Fig. S3). We also found that the total auto-fluorescence is proportional to the cell size (Fig. S3).

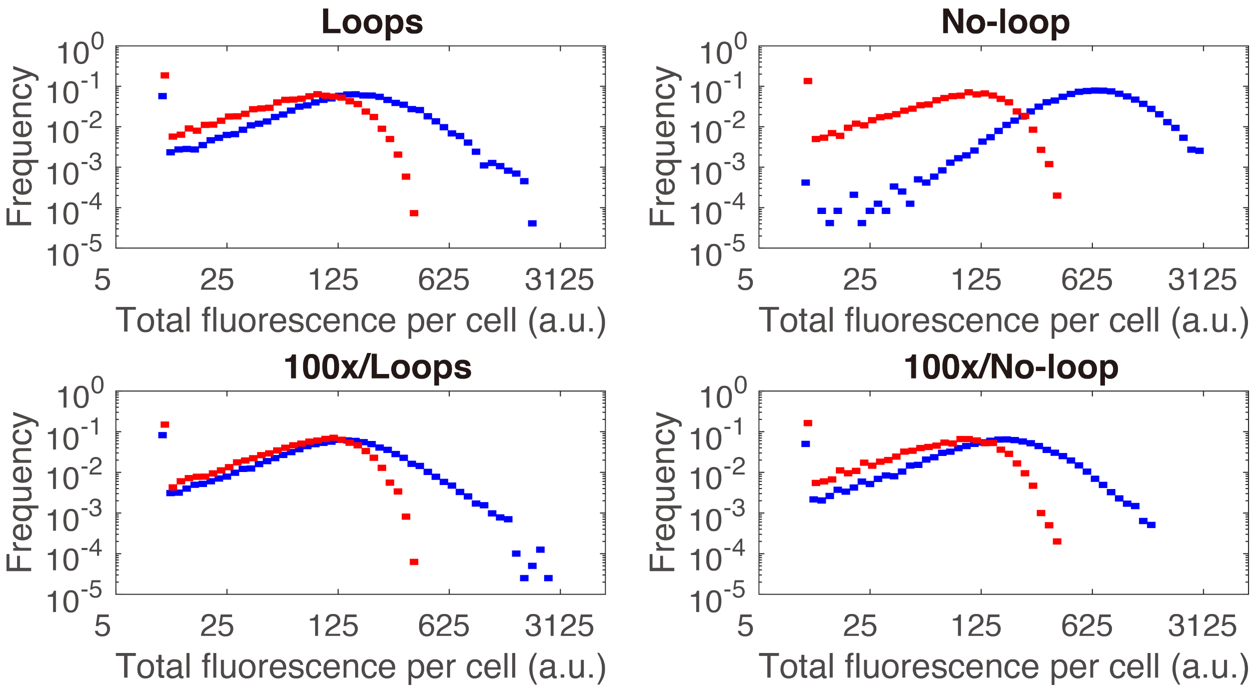

**Figure S2. Distributions of the total fluorescence per cell from the background strain and the four strains (Loops and No-loop) described in Table S1.** The red dots represent the distribution of the total auto-fluorescence per cell from the background strain CC-41 (table S1), and the blue dots represent the distribution of total fluorescence per cell for each of the four strains (Loops and No-loop) expressing Venus as described in table S1. The distribution from the Loops strain is almost identical to that of 100x/No-loop and also similar to 100x/Loop, but is different from that of the No-loop strain.

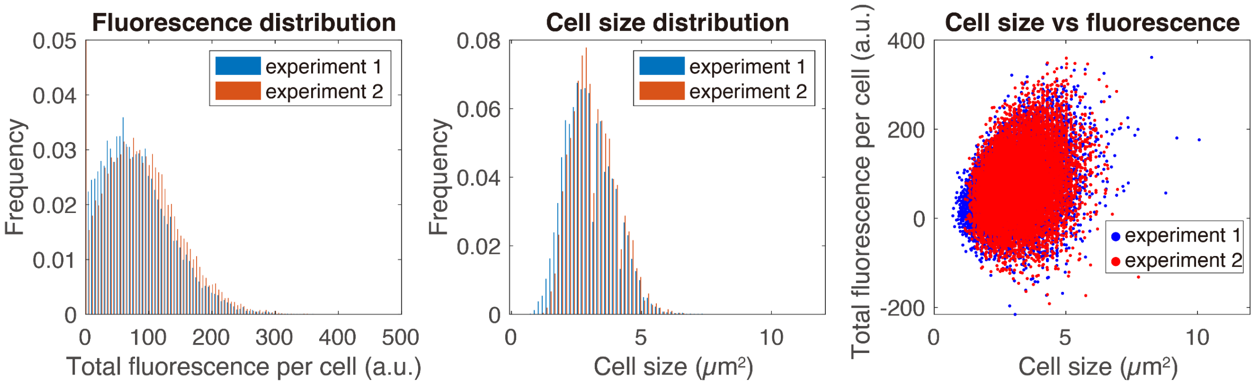

**Figure S3. Characterization of the auto-fluorescence signal from the background strain.** (left panel) The distributions of the total auto-fluorescence per cell, and (middle panel) the cell size distributions of background strain at different timepoints in two independent experiments. (right panel) Scatter plot of cell size versus total fluorescence per cell.

**Signal-to-noise ratio and the amplitude of pulses of the Loops and No-loop strains**

To determine to which extend we can distinguish the "true" signal from the background, we calculated the signal-to-noise ratio

$$SNR = \frac{\mu_{ON} - \mu_{background}}{\sigma_{background}} \quad (1)$$

where $\mu_{ON}$ is the mean fluorescent signal when the promoter is inferred as ON (see the later part of the Supplementary Information), and $\mu_{background}$ and $\sigma_{background}$ is the mean and the standard deviation of the auto-fluorescence calculated from the background strain that is grown within the same microfluidic chip as the Loops or No-loop strains. A Savitzky-Golay filtering (13) is applied to the fluorescence signals of both background and experimental strains. We also estimated the standard derivation for signal-to-noise ratio by applying Eq. (1) to all fluorescent signals when the promoter is ON. The signal-to-noise ratio is $6.0 \pm 5.2$ for the Loops strain, $5.9 \pm 5.2$ for the 100x/Loops strain, $6.0 \pm 4.7$ for the 100x/No-loop strain, and $18.0 \pm 10.5$ for the No-loop strain. Interestingly, the signal-to-noise ratio for the first three strains are similar.

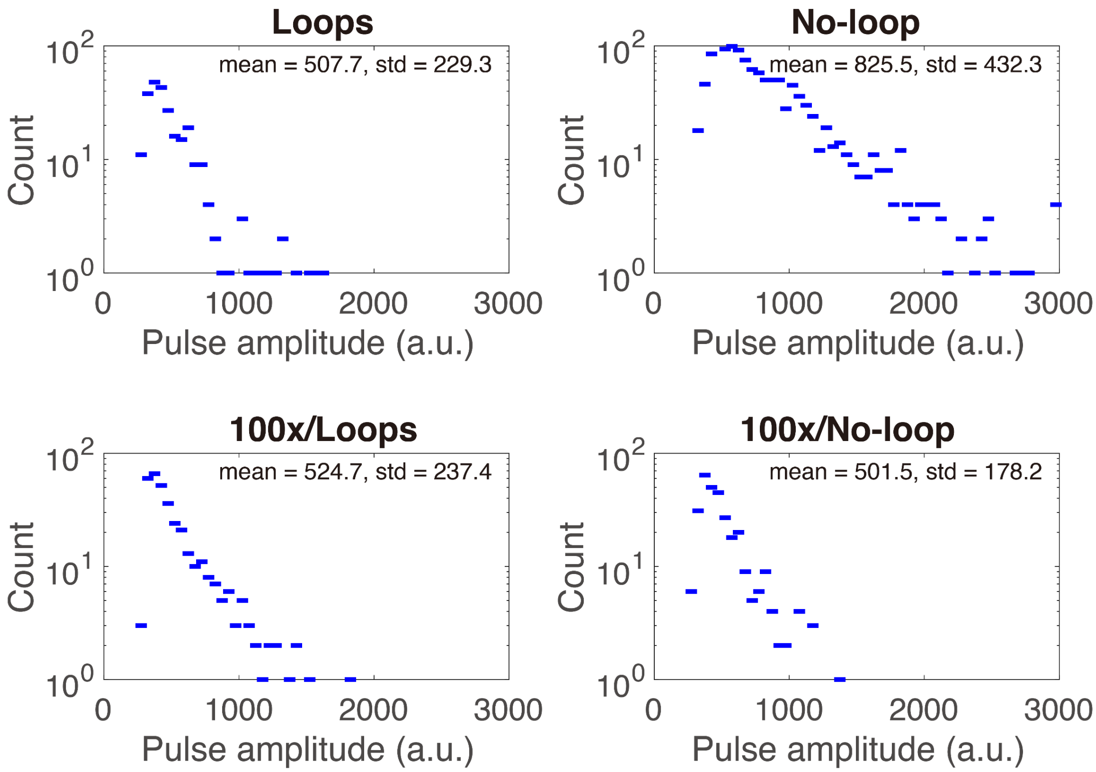

**Figure S4. Estimation of pulses amplitude.** To determine to which extend the "true" signal is separated from the background (shown in Fig. S3), we directly estimated the distributions of the pulse amplitude from fluorescence time series. In line with the observations from Fig. S2, the distribution of the Loops strain is similar to those of the 100x/Loops and 100x/No-loop strains, but different from that of the No-loop strain.

For the same purpose, we also make a rough estimate of the amplitude of the pulses observed from the Loops and No-loop strains. The estimate of pulse amplitudes is using only fluorescence signal within a single cell cycle (thus a pulse lasting more than one cell cycle will not be counted in this estimation). If the maximum fluorescence signal within a cell cycle is larger than a threshold (three sigma above mean background signals), and the difference between the maximum and the average fluorescence signals within the same cell cycle is larger than a second threshold (three times of background signals standard derivation), the difference between the maximum and the minimum fluorescence signals is considered as a candidate for estimating the pulses. We find that, the estimated amplitude of pulses from the loops strain is quite similar to those of the 100x/Loops and 100x/No-loop strains (Fig. S4). On the other hand, the estimated amplitude of the No-loop strain is larger than that of the Loops strain, but their pulses peaks are still of the same order (~ several hundred a.u.; Fig. S4). However, the ratio between the mean fluorescence level of the No-loop strain and the Loops strain is 3.97. Together, these observations suggest that the expression difference between the Loops strain and the No-loop strain is not solely caused by the amplitude of expression, the frequency of expression of the pulses may also play a role, since frequent protein expression can lead to higher fluorescence signal if the fluorescent proteins being produced are not diluted out when the second pulse happened.

**A probabilistic inference algorithm**

Intuitively, a 'hard threshold' method could be used for detecting the events of promoter expression. To define the state ‘ON’ of a promoter, one needs to calculate the change of the fluorescent signal between two frames, and compare with a predefined threshold.

In simulation, the 'hard threshold' method caused many false positive and false negative detections. We developed a probabilistic method for inferring the promoter activity, which outperforms the 'hard threshold' method in the regime of weak signals. This probabilistic method infers the promoter activity considering not only the cell-size dependent auto-fluorescence but also fluorescence signals of adjacent timepoints. We compute the maximum likelihood estimation to find out a set of non-negative promoter activity to minimize the weighted squared error between the observed signal and the sum of the background signal (that is a function of the cell size) and the underlying ‘real’ signal (a function of the promoter activity and the cell division).

*Likelihood function.* Let's denote *observed(t)* the total fluorescence per cell at time *t* for a lineage observed in an experiment. We use *ratio(t)* to denote the fraction of proteins that go to the next timepoint *t+1* that is a function of cell division,

$$ratio(t) = \begin{cases} 1.0, & \text{if cell does not divide at time t,} \\ <1.0, & \text{otherwise,} \end{cases} \quad (2)$$

*leakage(t)* to denote the promoter activity at time *t* that is the latent variable, which is always a non-negative number, and *real(t)* is used to denote the underlying 'real' signal as a function of *leakage(t)* and *ratio(t)*,

$$real(t) = real(t-1) * ratio(t-1) + leakage(t), \quad (3)$$

for $t \geq 2$. Since *ratio(t)* is given by the data, the information carried by *real(t)* and *leakage(t)* are equivalent. We further assume a normal distribution for the observation, i.e.,

$$observed(t) \sim N\big(real(t) + background(t), variance(t)\big), \quad (4)$$

where *background(t)* and *variance(t)* are the mean and variance of the auto-fluorescence per cell at time *t*, both of which can be directly calculated from the *cell size(t)* with the coefficients estimated from the total auto-fluorescence per cell of the background strain. The likelihood function could be written as

$$\begin{aligned} &\ln P(observed|real) \\ &= \sum_t \ln P\big(observed(t)\big|real(t)\big) \\ &= -\frac{1}{2}\sum_t \left( \ln variance(t) + \frac{\big(observed(t) - (real(t) + background(t))\big)^2}{variance(t)} \right) \\ &\qquad + \text{constant term,} \end{aligned}$$

by dropping off the first term in the parentheses that only depends on cell size, we have

$$-\sum_t \left( \frac{\big(observed(t) - (real(t) + background(t))\big)^2}{variance(t)} \right).$$

Thus, to maximize likelihood is equivalent to minimize the target function

$$\sum_t \left( \frac{\big(observed(t) - (real(t) + background(t))\big)^2}{variance(t)} \right), \quad (5)$$

$$\text{subject to } real(t) \geq 0,$$

a weighted sum of the difference between observed signal and 'real' signal plus background over all timepoints.

*Maximum likelihood solution: the estimation of leakage(t).* The maximum likelihood solution could be obtained by

$$\frac{\partial\ Eq.(5)}{\partial\ leakage(\tau)} = 0. \tag{6}$$

The only term in Eq. (5) that depends on $leakage(\tau)$ is *real(t)* for timepoints $t \geq \tau$. As shown in Eq. (3), the dependence of *real(t) on* $leakage(\tau)$ has direction in time. Let's denote the timepoints of cell divisions $\omega_1, \omega_2 \dots, \omega_m$ between *t* and $\tau$, where *m* is short for $m(t, \tau)$. Thus, for *real(t)* with $t \geq \tau$,

$$\begin{aligned} real(t) &= real(t-1) * ratio(t-1) + leakage(t) \\ &= real(\omega_m + 1) + \sum_{\omega=\omega_m+1}^{t} leakage(\omega) \\ &= ratio(\omega_m) \times real(\omega_{m-1} + 1) + ratio(\omega_m) \times \sum_{\omega=\omega_{m-1}+1}^{\omega_m} leakage(\omega) \\ &\quad + \sum_{\omega=\omega_m+1}^{t} leakage(\omega) \\ &= ratio(\omega_{m-1}) \times ratio(\omega_m) \times real(\omega_{m-2} + 1) \\ &\quad + ratio(\omega_{m-1}) \times ratio(\omega_m) \times \sum_{\omega=\omega_{m-2}+1}^{\omega_{m-1}} leakage(\omega) \\ &\quad + ratio(\omega_m) \times \sum_{\omega=\omega_{m-1}+1}^{\omega_m} leakage(\omega) + \sum_{\omega=\omega_m+1}^{t} leakage(\omega) \\ &= \prod_{i=1}^{m} ratio(\omega_i) \times real(\tau) + effective\ leakage(t, \tau) \end{aligned} \tag{7}$$

with

$$effective\ leakage(t, \tau) = \sum_{j=0}^{m-1} \prod_{i=m-j}^{m} ratio(\omega_i) \sum_{\omega=\omega_{m-j-1}+1}^{\omega_{m-j}} leakage(\omega) + \sum_{\omega=\omega_m+1}^{t} leakage(\omega), \tag{8}$$

by denoting $\omega_0 = \tau$. We further denote $t_{end}$ as the last timepoint. Thus, starting from Eq. (6),

$$\frac{\partial\, Eq.(5)}{\partial\, leakage(\tau)} = \frac{\partial \sum_{t=\tau}^{t_{end}} \left( \frac{\left(observed(t) - \left(real(t) + background(t)\right)\right)^2}{variance(t)} \right)}{\partial\, leakage(\tau)}$$

$$= \sum_{t=\tau}^{t_{end}} \frac{\partial \left( \frac{\left(observed(t) - \left(real(t) + background(t)\right)\right)^2}{variance(t)} \right)}{\partial\, leakage(\tau)}$$

$$= -2 \sum_{t=\tau}^{t_{end}} \frac{observed(t) - \left(real(t) + background(t)\right)}{variance(t)} \frac{\partial\, real(t)}{\partial\, leakage(\tau)}$$

$$= -2 \sum_{t=\tau}^{t_{end}} \frac{observed(t) - \left(real(t) + background(t)\right)}{variance(t)} \prod_{i=1}^{m(t,\tau)} ratio(\omega_i) \qquad (9)$$

$$= 0.$$

By inserting Eq. (7) into Eq. (9), we have

$$\sum_{t=\tau}^{t_{end}} \frac{observed(t) - \left(\prod_{i=1}^{m(t,\tau)} ratio(\omega_i) \times real(\tau) + effective\ leakage(t,\tau) + background(t)\right)}{variance(t)}$$

$$\times \prod_{i=1}^{m(t,\tau)} ratio(\omega_i) = 0$$

$$\sum_{t=\tau}^{t_{end}} \frac{observed(t) - \left(effective\ leakage(t,\tau) + background(t)\right)}{variance(t)} \times \prod_{i=1}^{m(t,\tau)} ratio(\omega_i)$$

$$= \sum_{t=\tau}^{t_{end}} \frac{real(\tau)}{variance(t)} \times \left( \prod_{i=1}^{m(t,\tau)} ratio(\omega_i) \right)^2$$

we have

$$real(\tau) = \frac{\sum_{t=\tau}^{t_{end}} \frac{observed(t) - \left(effective\ leakage(t,\tau) + background(t)\right)}{variance(t)} \times \prod_{i=1}^{m(t,\tau)} ratio(\omega_i)}{\sum_{t=\tau}^{t_{end}} \frac{1}{variance(t)} \times \left(\prod_{i=1}^{m(t,\tau)} ratio(\omega_i)\right)^2}, \qquad (10)$$

the physical meaning of which is a weighted sum of the difference between observation and background plus the effective leakage estimated in current iteration over timepoints after $\tau$. $leakage(\tau)$ could be easily calculated by combining Eq. (3) and Eq. (10):

$$leakage(\tau) = real(\tau) - real(\tau - 1) * ratio(\tau - 1), \tag{11}$$

for $\tau \geq 2$, and $leakage(\tau) = real(\tau)$ for $\tau = 1$, which is the leakage before observation.

*Estimation of background(t) and variance(t).* From the data of the background strain, we find that both the mean and the standard derivation of the total auto-fluorescence per cell are linearly proportional to the cell size. We can directly estimate the dependence of *background(t)* to *cell size(t)* using a linear model. On the other hand, we estimate *variance(t)* from *cell size(t)* based on the squared error between the observed signal and the background signal estimated using the linear model,

$$squared\ error(t) = \left(observed(t) - background(t)\right)^2. \tag{12}$$

The squared error could be decomposed into two parts: the cell size-dependent variance with coefficients $\rho_{cell\ size\ squared}$ and $\rho_{cell\ size}$, and the part of variance that is directly from the observation with coefficient $\rho_{observation}$. We have

$$\left(\overrightarrow{cell\ size^2}, \overrightarrow{cell\ size}, \vec{1}\right)\begin{pmatrix}\rho_{cell\ size\ squared}\\ \rho_{cell\ size}\\ \rho_{observation}\end{pmatrix} = \overrightarrow{squared\ error}. \tag{13}$$

Thus, the least squared solution for those coefficients could be calculated by

$$\begin{pmatrix}\rho_{cell\ size\ squared}\\ \rho_{cell\ size}\\ \rho_{observation}\end{pmatrix} = \overrightarrow{squared\ error} \times \left(\overrightarrow{cell\ size^2}, \overrightarrow{cell\ size}, \vec{1}\right)^{-1}. \tag{14}$$

*Protein partition: the estimation of ratio(t) for a cell division*. The fraction of the 'real' signal that a daughter cell inherits from the mother cell is directly estimated by the ratio between their total fluorescence per cell, i.e., if a cell divides at timepoint *t-1*, we have

$$ratio(t) = \frac{observed(t-1)}{observed(t)}, \tag{15}$$

with a lower boundary set to be 0.01 and an upper boundary set to be 0.99.

*Parameter updating rule to find out the local maxima of likelihood.* The overall pipeline for the algorithm is available at Fig. S5a. To infer promoter activity from the time series data, we run ten rounds of fitting and select the one with largest likelihood score as the final result. In each round of the fitting, we obtain the initial values of the promoter activity without the nonnegative restriction, and update their values with the nonnegative restriction (see *Nonnegative restriction of promoter activity* in Supplementary Information). Based on the promoter activity estimated with the nonnegative restriction, we further update their values in a 'diffusion' way. For each pulse, we checked if the likelihood could be improved by 'diffusing' a part of the promoter activity to its adjacent timepoints. These steps are repeated five times in each round of fitting, and the updated parameters from the precedent run is used as the initial parameters for the subsequent run. The pseudocode for finding out the maximum likelihood is available in Fig. S5b.

(a) **Overall pipeline for the algorithm**

(1) Estimate $\rho_{\text{cell size squared}}$, $\rho_{\text{cell size}}$ and $\rho_{\text{observation}}$ from the background strain using Eq. (14).

(2) Infer the promoter activity for each lineage of a strain.

(2.a) obtain initial parameters without nonnegative restriction using panel *b*;

(2.b) update parameters with nonnegative restriction using panel *b*;

(2.c) update parameters for each pulse in a 'diffusion' way.

(3) Calculate and merge the statistics of a strain from all its lineages.

(b) **Pseudo-code for inferring promoter activity**

```
for fitting_round = 1 to 10
   for run_round = 1 to 5
      update leakage(t=1) using panel c;
      for τ = random_shuffle(2, t_end)
         update leakage(t= τ) using panel c;
      end
   end
   calculate likelihood(fitting_round) using Eq. (5);
end
return the parameters with maximum likelihood;
```

(c) **Pseudo-code for updating leakage(t)**

```
if observed(t) > real(t - 1) + background(t) + ρ_signal * background_std(t) and
   leakage(t) > μ(background_derivative) + ρ_derivative * σ(background_derivative)
     leakage(t) = Eq. (11) if without nonnegative restriction,
                = max(0.0, Eq. (11)) if with nonnegative restriction.
elseif with nonnegative restriction
     leakage(t) = 0.0;
end
```

**Figure S5. Pseudo-code for our pulse detection algorithm.** (a) Overall pipeline for the algorithm. (b) Pseudo-code for inferring promoter activity. The function *random_shuffle* returns a permutation for the input number array in which elements are randomly rearranged. (c) Pseudo-code for updating *leakage(t)*. *leakage(t)*, *observed(t)*, *real(t)*, *background(t)*, and *variance(t)* are defined in the 'a probabilistic inference algorithm' section of the supplementary information. *background_std(t)* is the squared root of the *variance(t)*. $\mu$(background_derivative) and $\sigma$(background_derivative) are the mean and the standard derivation of the first-order derivative of the total auto-fluorescence per cell calculated from the background strain. The function *max* returns the maximum value of the numbers.

*Nonnegative restriction of promoter activity.* Parameter updating to search the local maxima with the constraint of non-negative promoter activity is challenging. To solve this problem, we update the parameters in two rounds. In the first round, we update the parameters without the nonnegative restriction, so that the sum of the background signal and the 'real' signal would match the observed signal at all timepoints for a lineage. In the second round, the nonnegative promoter activity constraint is introduced using the parameters estimated from the first round as the starting point of iteration. Detailly, with or without the nonnegative restriction, the first step is to update the promoter activity of the first timepoint in order to estimate the amount of fluorescence signals for proteins being produced before the start of the observation. In the second step, we update the promoter activity for each of the remaining timepoints in a randomized order to prevent visiting the same local likelihood maxima in different realizations.

*Thresholds for parameter updating at each timepoint*. For the parameter updating at each timepoint, we accept the update of the promoter activity only if it passes a given threshold; otherwise, this promoter activity will be set to be zero. The threshold we set would not only depends on the total fluorescence per cell at the current timepoint, but also the temporal first-order derivative of the total fluorescence per cell (Fig. S5c).

Depending on the levels of total fluorescence per cell and burst size, we use two sets of thresholds for strains in our study. For the Loops, 100x/Loops, 100x/$O_1$-$O_3$ One-loop, $O_1$-$O_2$ One-loop, 100x/$O_1$-$O_2$ One-loop, and 100x/No-loop strains, $\rho_{signal} = 2.5$, and $\rho_{derivative} = 1.5$, which is a stringent threshold. For the $O_1$-$O_3$ One-loop and No-loop strains, a loose threshold is used. We set $\rho_{signal} = 2.5$, and there is no threshold in fluorescence derivative.

*Data preprocessing.* We apply a Savitzky-Golay filtering for data smoothing within each cell cycle for the input time series data before inference.

*Statistics of expression dynamics*. The intervals of promoter activity corresponding to ON and OFF states that could be calculated from the inferred promoter activity using this probabilistic method. An interval 'ON' is defined as the duration of contiguous timepoints with nonzero promoter activity, and an interval of 'OFF' activity is the duration between two ON intervals. An ON/OFF interval at the beginning/end of the time series whose duration cannot be accurately determined, however, will be ignored. In our experiments, the duration of an ON interval is usually shorter than one cell cycle. In power spectrum analyses (Figs. S6-S9 (a), lower panels), the power of the fluorescence signals of the background strain in the high frequency domain (< a cell cycle) is higher than that of a Loops or No-loop strain, making it unrealistic to reliably detect ON intervals. The burst size of a pulse associated with an ON interval, which is defined as the integral of the promoter activity, is a more robust measurement. Thus, we will use OFF intervals and burst size to characterize the promoter activity. We also found that a longer ON interval sometimes will be detected as two or more shorter ON intervals with small gaps. Thus, for our probabilistic method, 'ON' intervals with gaps smaller or equal to 15 minutes, as well as the associated burst size, will be merged, which would largely improve the performance of simulation mentioned in the later part of the Supplementary Information.

**Simulation of artificial time series to evaluate error rates in pulse detection using our probabilistic method**

Experimental fluorescence time series from the background strain is used as the starting point for the simulations. There are several controlling parameters that are used to adjust the statistics of the simulated time series so that they resemble the time series observed in our experiments: the parameters called *OFF interval* and *ON interval* control the duration of an OFF or ON interval, and the parameter called *burst size* controls the burst size of a pulse, which is randomly partitioned into the promoter activity of each timepoint of an ON interval. Motivated by Fig. 3 in the main text, we have an additional parameter controlling the ratio of the mean promoter activity between the first and the second half of a cell cycle. The promoter activity at the second half of a cell cycle would be scaled with that ratio to represent the correlation between promoter activity and cell cycle. Each controlling parameter could be a constant number or random numbers drawn from a distribution. Once the promoter activity of all timepoints is generated, it is added to a real time series associated with the auto-fluorescence of the background strain. A daughter cell would inherit only a fraction of the added signals from the mother cell.

The performance of the algorithm depends on the values chosen for the controlling parameters of the pulses. To evaluate the error rate in pulse detection, it is crucial to identify a proper parameter set for which not only the distribution of the total fluorescence per cell and the distribution of the fluorescence derivative of the simulations and experiments match, but also their power spectra.

The search of the parameters is performed manually. The parameters of the Loops strain and the 100x/No-loop strain are the same (i.e., they share the same set of functions for generating random variables), the duration of an OFF interval is on average 220 minutes, with a burst size that is 2.71 fold the mean fluorescence signals of the background and with a duration distributed around 25 min (Figs. S6 and S10). The parameters for the 100x/Loops strain are identical to that of the Loops strain but an OFF interval, however, lasts on average 395 minutes (Fig. S7). On the other hand, for the parameters of the No-loop strain, the duration of an OFF interval is distributed around 35 minutes, the burst size is 3.69 fold of the mean fluorescence signals of the background and the ON duration is distributed around 38 minutes (Fig. S8). For all Loops and No-loop strains, the ratio between the first and the second half of the cell cycle is set to be 1.7 (motivated by the ratio from the experiments; see Fig. S14).

As shown in Figs. S6-S10, for the simulations of the Loops and No-loop strains, overall the distributions of the detected OFF intervals agree with the real distributions. The upper boundaries of OFF intervals in inference also agree with the real situation, with occasional false positives being introduced at relatively low frequency. Our pulse detection algorithm can also overall accurately detect the distributions of the burst size for the pulses except those with very small burst size (Figs. S6-S10).

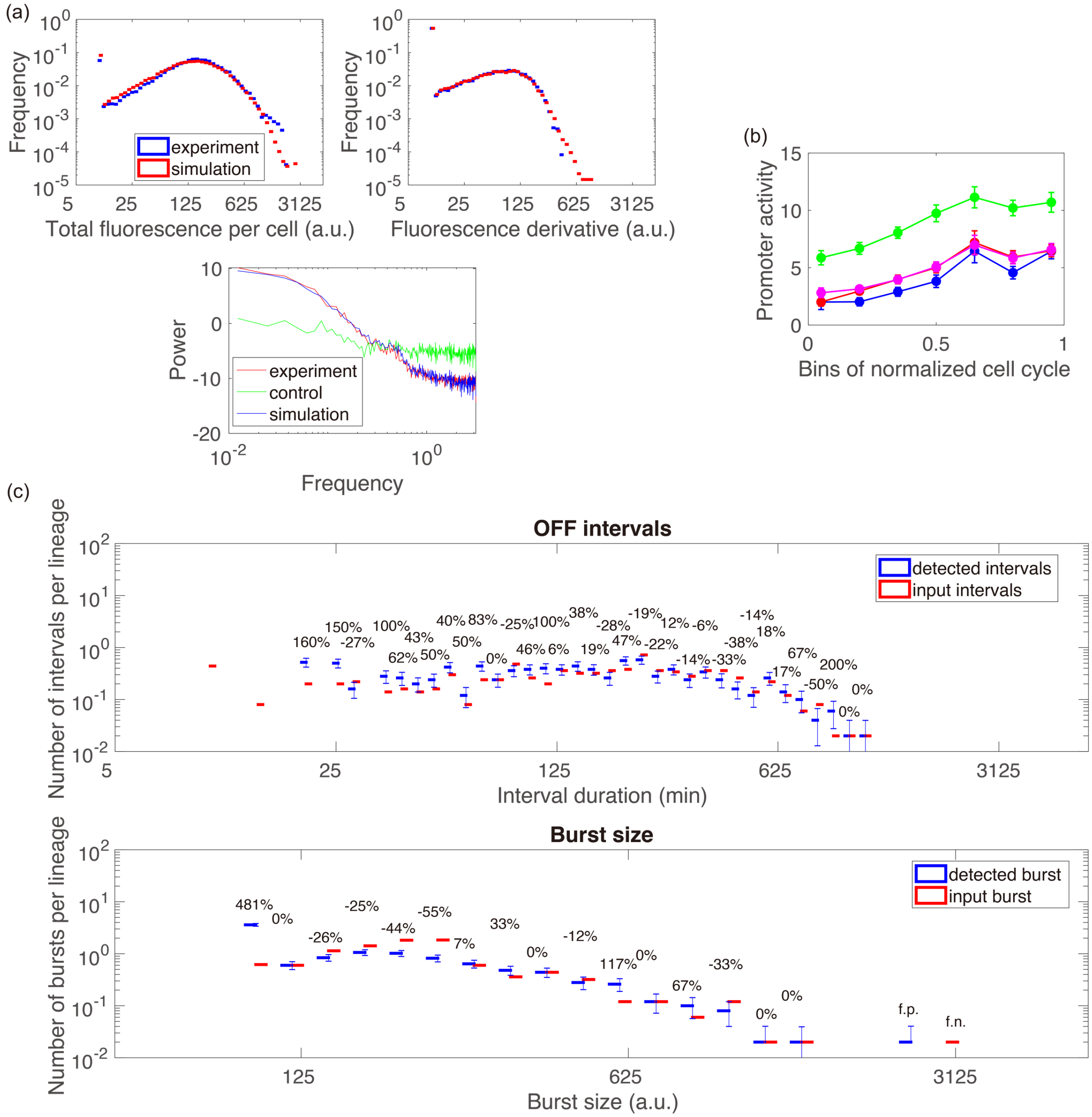

**Figure S6. Parameters and simulation tests for the Loops strain.** (a) The goal is to simulate time series that have similar distribution of the total fluorescence per cell, distribution of fluorescence derivative, and power spectrum as for the time series measured directly from growing bacteria. In our simulation, the *OFF interval* and *ON interval* is a multiple of 5 minute dwell time, and the *burst size* is measured in term of number of folds of the mean fluorescence of the background strain. We chose for the parameters of the Loops strain as follow. *OFF interval*: geornd(1/45.0); *ON interval*: 3 + geornd(1/3.0); *burst size*: max(2.0, geornd(1/2.75))

(mean=2.71, determined by numeric). The function *max* returns the maximum value of the numbers, function *round* returns the nearest integer to a float number, and function *georndd* gives a random number from the geometric distribution (p.m.f. $f(x) = p(1-p)^x, x = 0,1,2,\ldots$), respectively. (b) The correlation between promoter activity and cell cycle inferred by the algorithm (blue line) matches that from the input (the red line represents the input leakage, and the purple line is the input leakage after applying a Savitzky-Golay filter). The fluorescence derivative that is directly calculated from the data, however, overestimate the level of the promoter activity. An error bar represents the coefficient of variation. (c) The upper panel gives the distributions of input and detected OFF interval duration, and the lower panel shows the distributions of input and detected burst size. Y-axis gives the number of OFF intervals or pulses with a burst size per lineage for a given bin. The error bar of each bin is the standard derivation generated using the bootstrapping method. f.p. (i.e. false positive) means that there is actually no OFF interval or burst size in the associated bin and the inference yields a false positive, and f.n. (i.e. false negative) means that there is actually an OFF interval or a pulse with a burst size but it has not been detected.

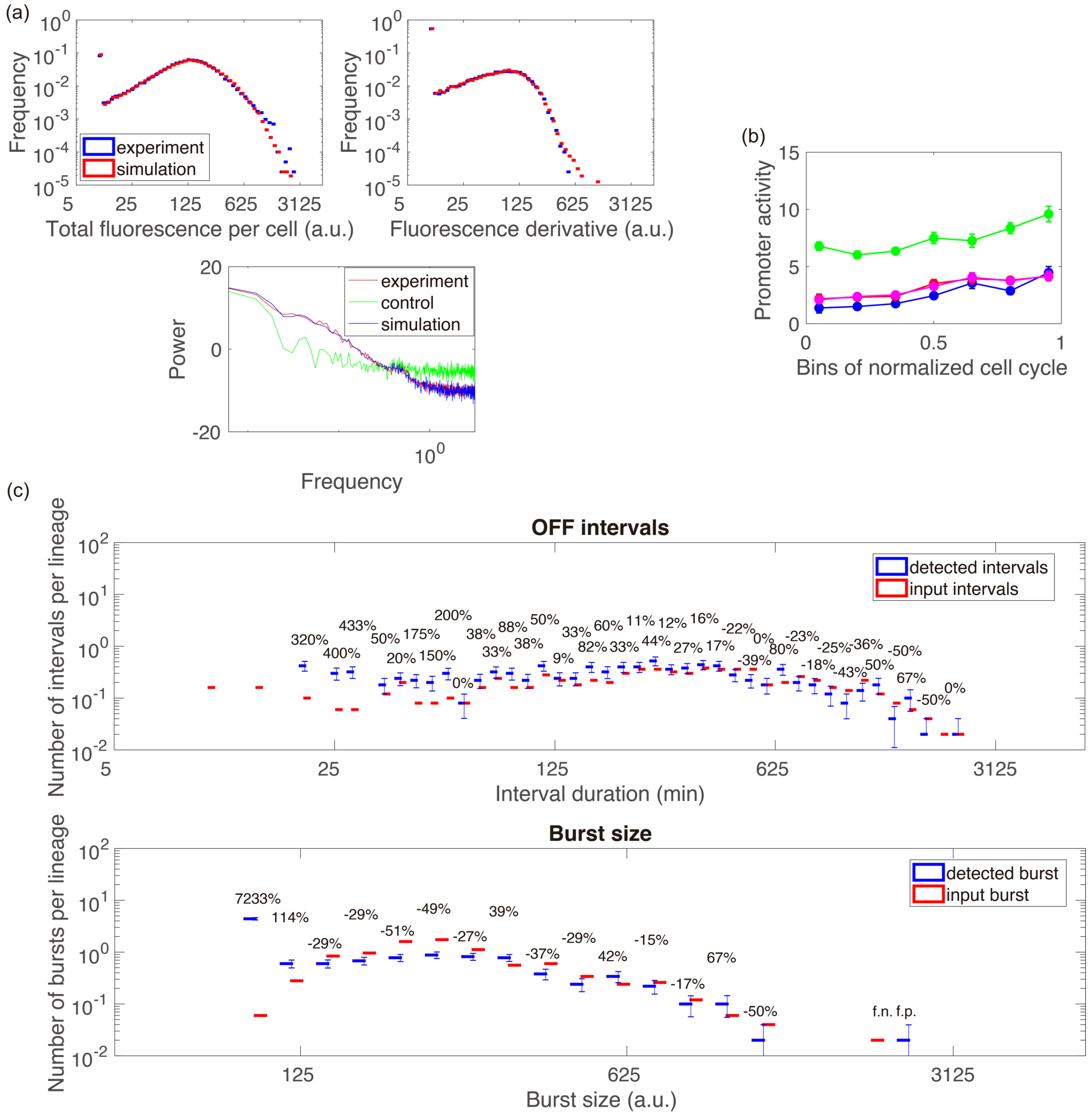

**Figure S7. Parameters and simulation tests for the 100x/Loops strain.** The parameters of 100x/Loops: *OFF interval*: geornd(1/80.0); *ON interval*: 3 + geornd(1/3.0)); *burst size*: max(2.0, geornd(1/2.75)). The meaning of the symbols of this figure is identical to that of Fig. S6.

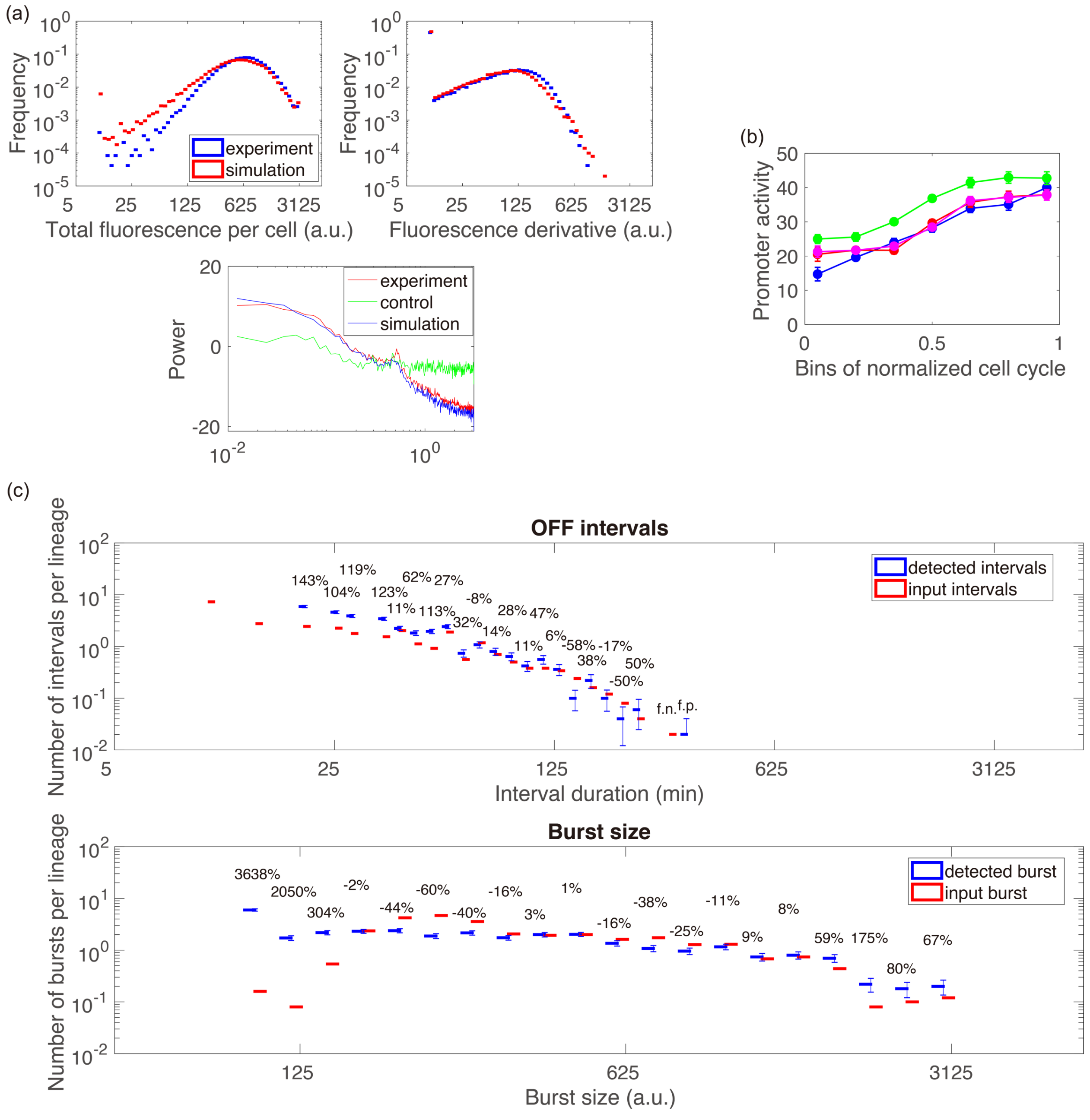

**Figure S8. Parameters and simulation tests for the No-loop strain.** The parameters of the No-loop strain: *OFF interval*: geornd(1/8.0); *ON interval*: 4 + geornd(1/4.5); *burst size*: max(2.0, geornd(1/4.0)) (mean=3.69, determined by numeric). The meaning of the symbols of this figure is identical to that of Fig. S6.

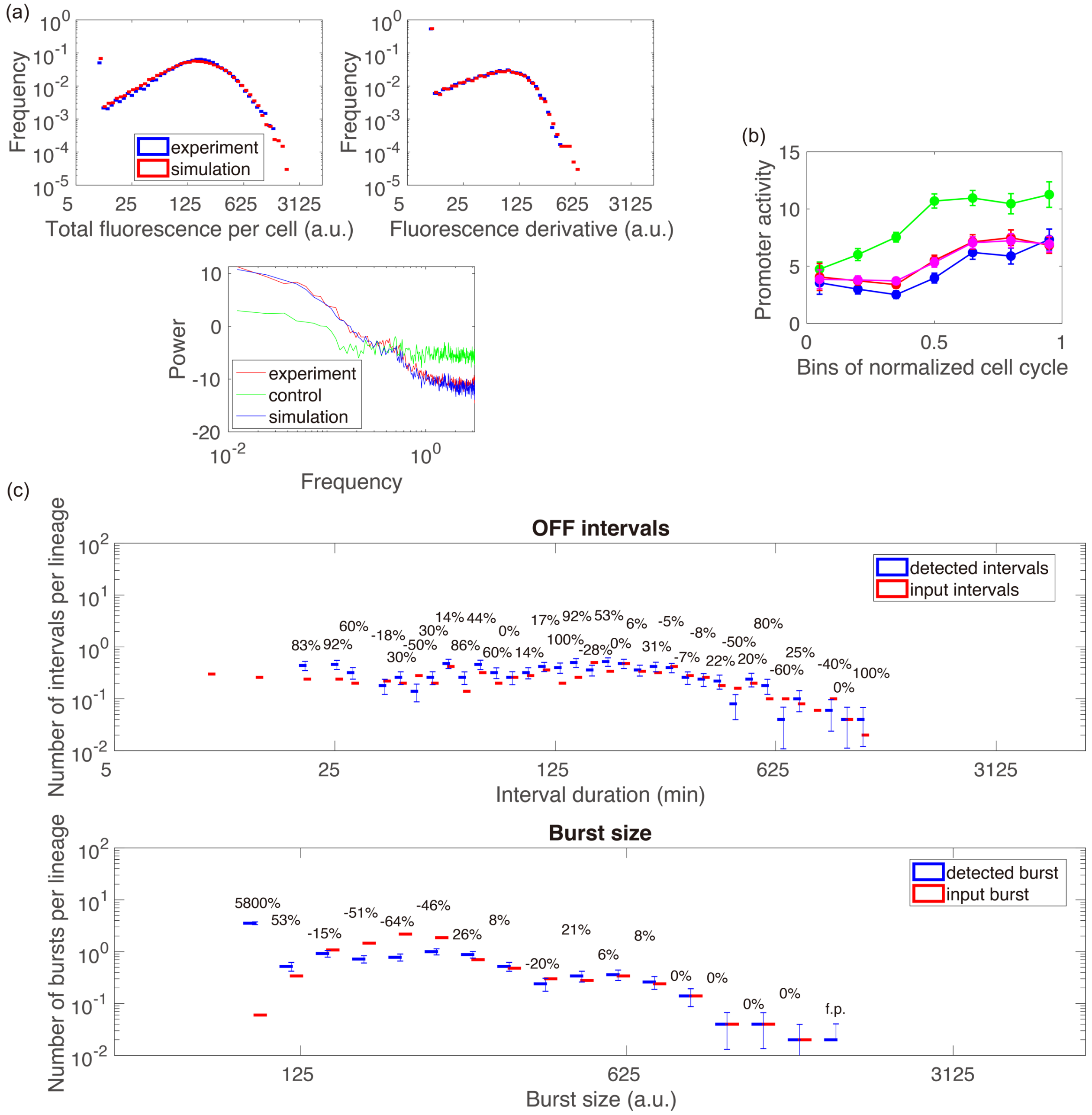

**Figure S9. Parameters and simulation tests for the 100x/No-loop strain.** The parameters of 100x/No-loop is identical to those of the Loops strain, but the experimental fluorescence time series of the background strain in simulation is from the same experiment of 100x/No-loop. The meaning of the symbols of this figure is identical to that of Fig. S6.

**‘Scaling’ test for simulated time series**

We performed a ‘scaling’ test to check whether long OFF intervals observed in the time series of the Loops strain, the 100x/Loops strain and the 100x/No-loop strain could be merely explained by the difference in amplitude comparing to the No-loop strain, as the pulse amplitude underlies the detectability of the OFF intervals. For example, extremely weak amplitude could lead to false positive and yield longer OFF intervals.

In our simulations, the burst size of the pulses for the Loops strain, the 100x/Loops strain and the 100x/No-loop strains is 2.71 fold the mean fluorescence signals of the background strain, but for the No-loop strain is 3.69 fold. We scaled the amplitude of the Loops, 100x/Loops and 100x/No-loop strains to that of the No-loop strain, and *vice versa* and check the effect of these different scaling on the detection of the longest OFF intervals. The detected OFF interval distributions of the ‘scaling’ test for all strains are close to the real distributions (Fig. S10). Therefore, the presence of long OFF intervals observed for the three strains (Loops, 100x/Loops and 100x/No-loop) cannot be explained by error in pulses detection due to the weak amplitude of the pulses.

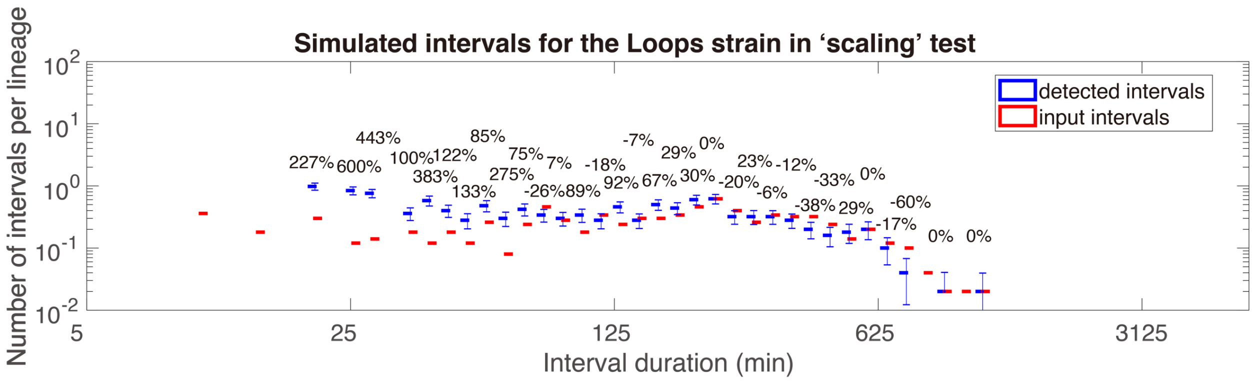
Simulated intervals for the Loops strain in ‘scaling’ test
detected intervals
input intervals
Number of intervals per lineage
Interval duration (min)

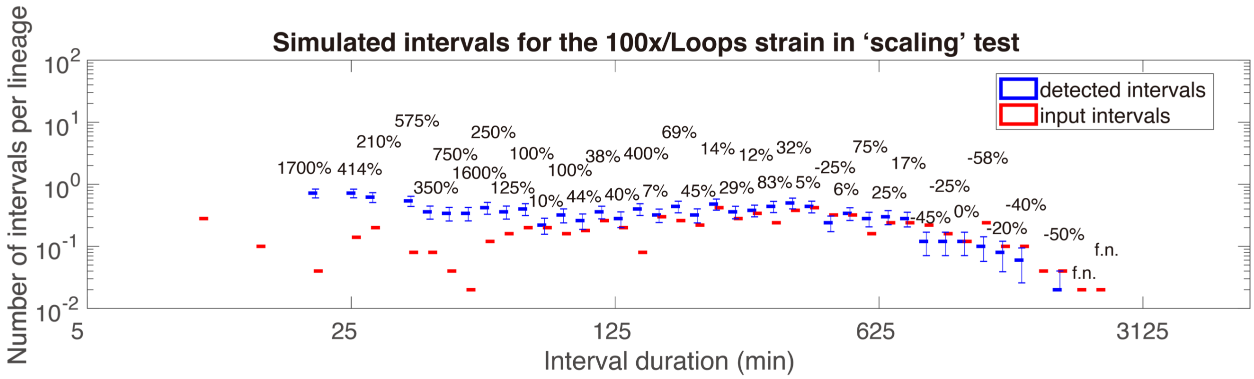
Simulated intervals for the 100x/Loops strain in ‘scaling’ test
detected intervals
input intervals
Number of intervals per lineage
Interval duration (min)

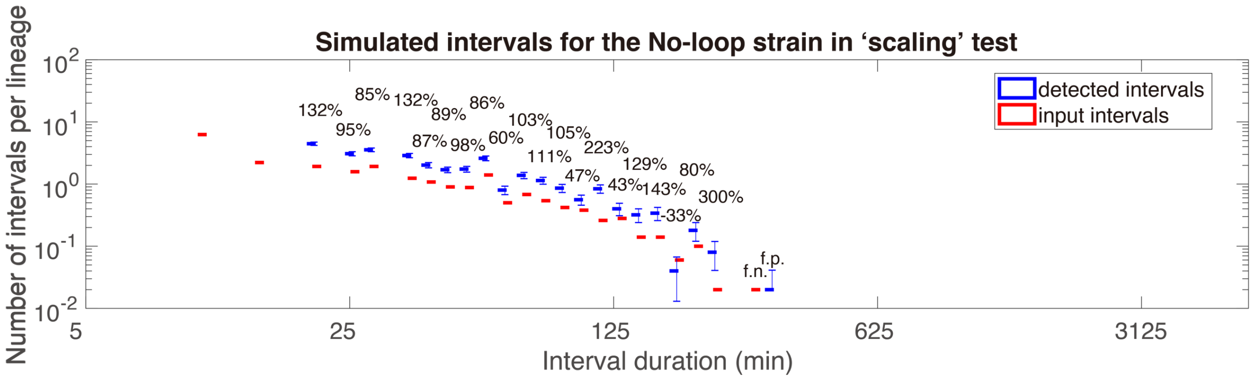
Simulated intervals for the No-loop strain in ‘scaling’ test
detected intervals
input intervals
Number of intervals per lineage
Interval duration (min)

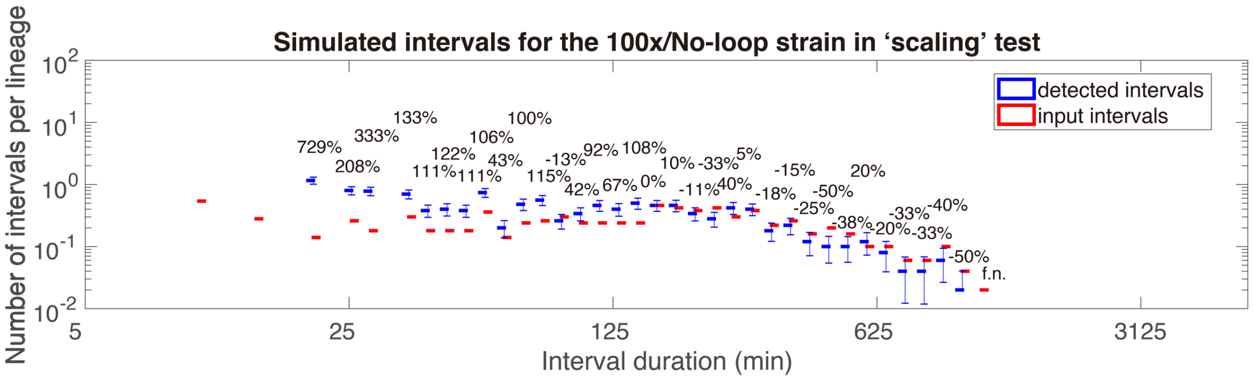
Simulated intervals for the 100x/No-loop strain in ‘scaling’ test
detected intervals
input intervals
Number of intervals per lineage
Interval duration (min)

**Figure S10. 'Scaling' test for the simulation mentioned in Fig. S7**. For each simulated time series generated in Figs. S6-S9, we performed a 'scaling' test. This test consisted in scaling the magnitude of burst size with a certain factor. Specifically, the burst size of the Loops, 100x/Loops and 100x/No-loop strains is scaled with a factor of 1.36, and that of the No-loop strain is scaled with 0.74, so that the burst size of the Loops, 100x/Loops and 100x/No-loop strains are scaled to that of the No-loop strain, and *vice versa*. The meaning of the symbols of this figure is identical to that of Fig. S6c.

**Inference using a 'hard threshold' method**

we compared the 'hard threshold' method to our probabilistic pulse detection algorithm, using the same simulated data set as in Figs. S6-S9. The optimal 'hard threshold' is selected to minimize the average Kullback–Leibler divergence between the input distributions of OFF intervals and those of the detected OFF intervals using various combinations of pulse size and frequency. Similar to the probabilistic detection algorithm, a Savitzky-Golay filtering is used in the data pre-processing for the 'hard threshold' method. Overall, the 'hard threshold' method yields larger differences between the input and detected distributions for not only OFF intervals but also burst size comparing to those the probabilistic algorithm (Figs. S6-S9 vs. Figs. S11-S12).

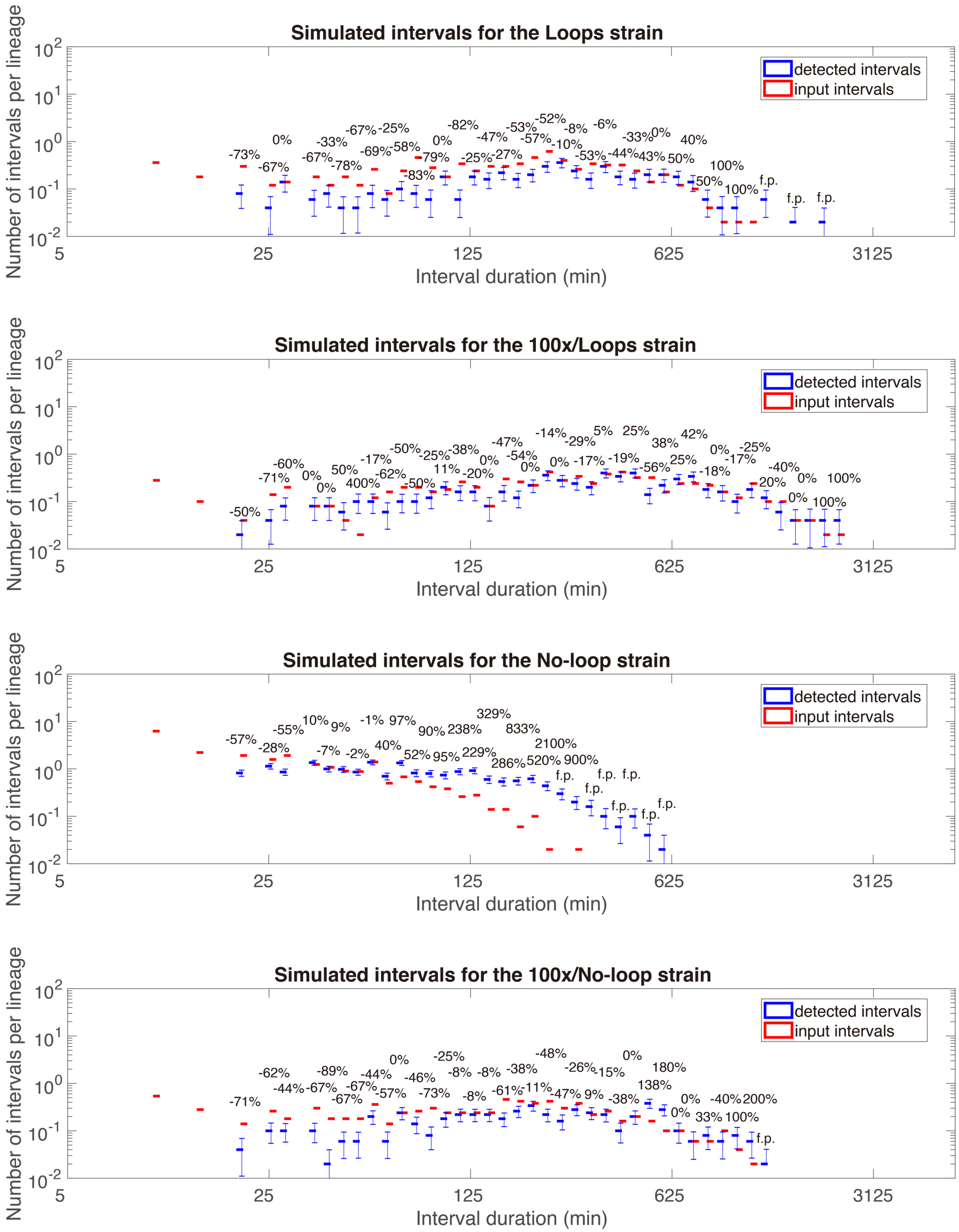

**Figure S11. Simulated tests for OFF intervals detection using 'hard threshold' method.** In order to compare the 'hard threshold' method with our pulse detection algorithm, we use the former to detect the OFF intervals using the same simulated data set to Figs. S6-S9. The distributions of input and actual OFF intervals are given for simulation of each strain. The meaning of the symbols of this figure is identical to that of Fig. S6c.

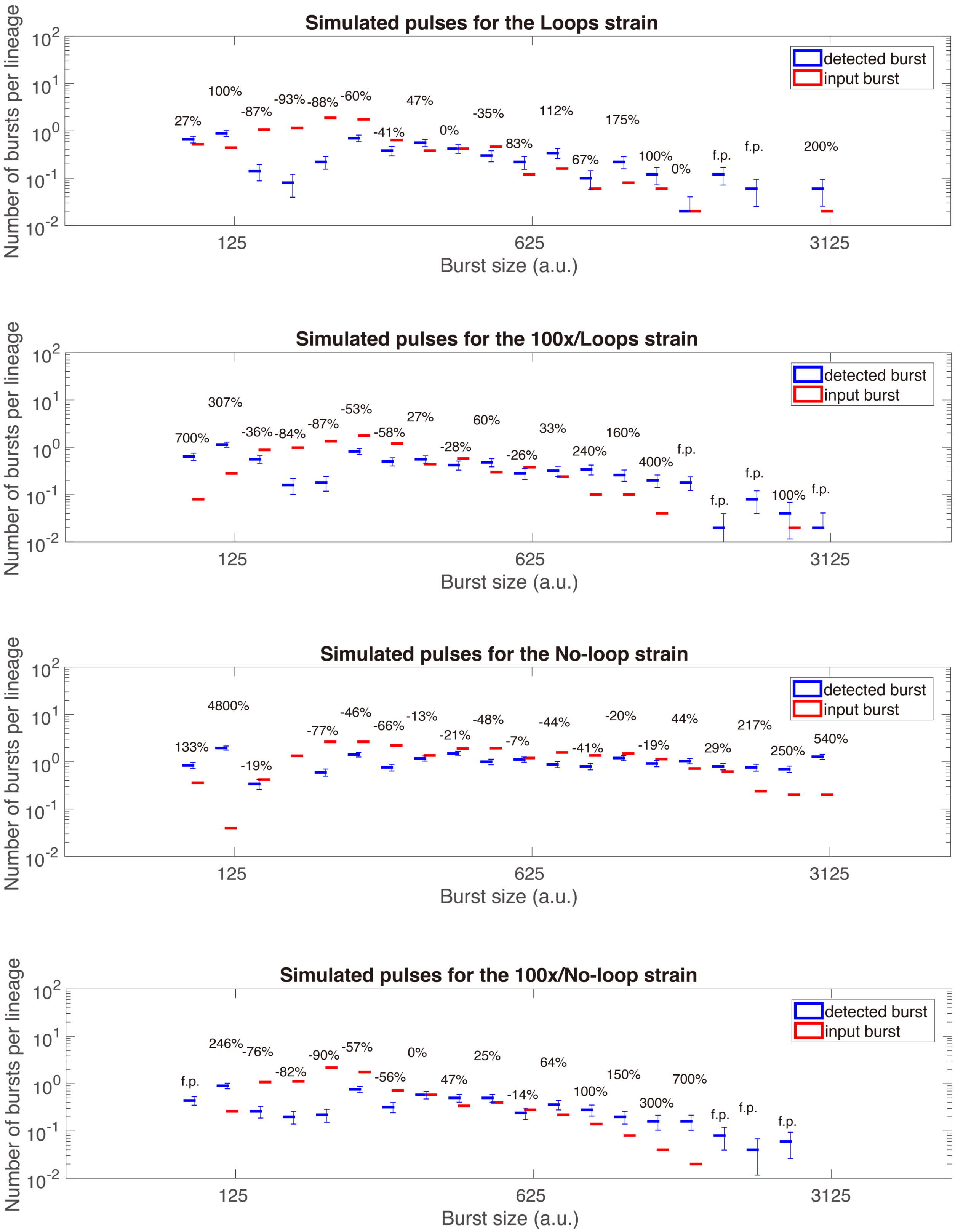

**Figure S12. Simulated tests for burst size detection using hard threshold method.** The distributions of input and detected burst size of pulses associated with Fig. S11 are given. The meaning of the symbols of this figure is identical to that of Fig. S6c.

**OFF interval distributions for One-loop strains**

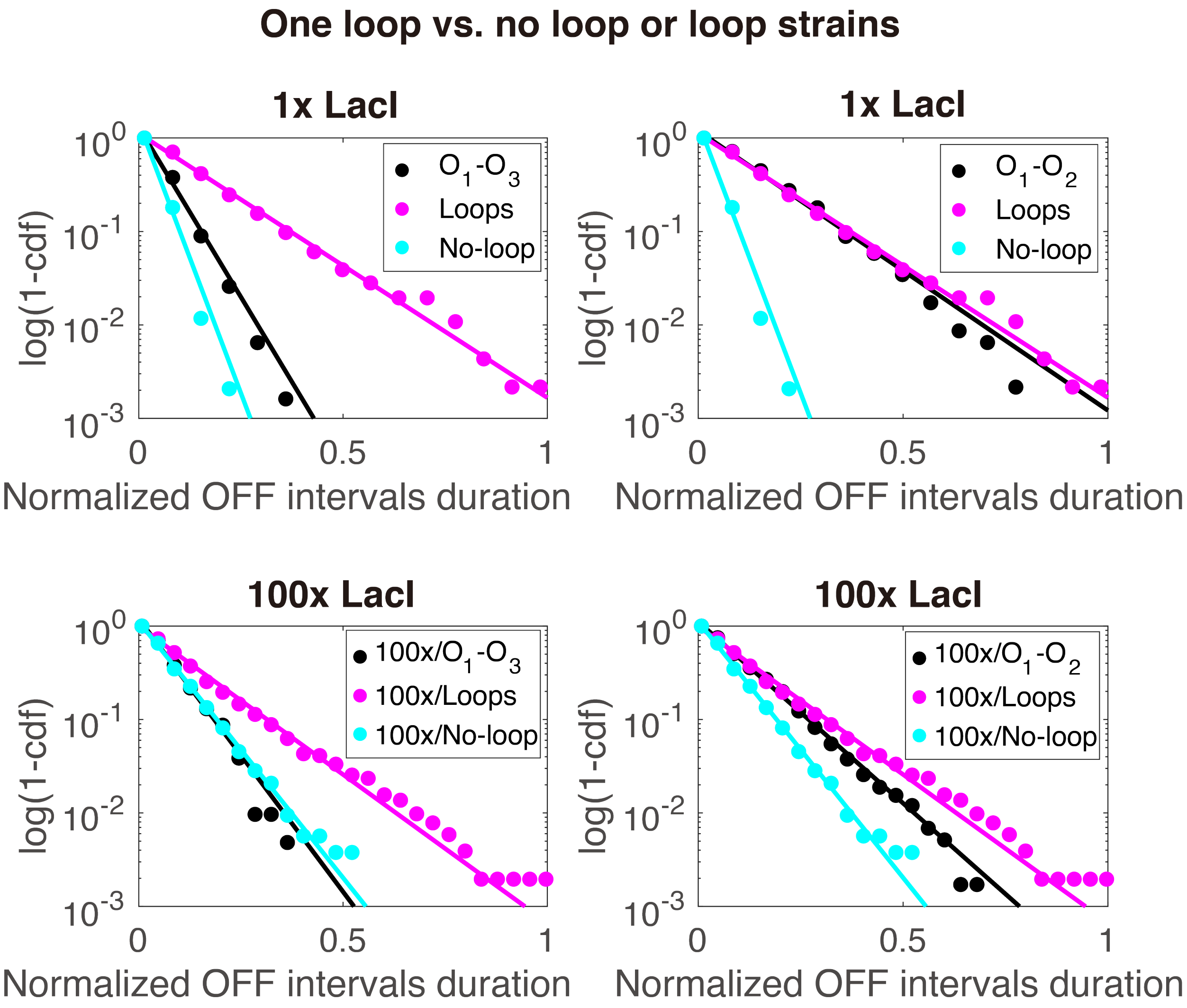

**Fig. S13. OFF intervals for One-loop strains follow exponential distributions.** The cumulative distributions ($P(X \geq x)$) for four One-loop strains were compared with those of the Loops and No-loop strains. For each panel, the OFF intervals were normalized by dividing the maximum value from magenta dots. Normalization factors: upper panels (1440 min), and lower panels (2520 min). Slopes before normalization: $O_1$-$O_3$ One-loop (-0.01162 $min^{-1}$), $O_1$-$O_2$ One-loop (-0.00478 $min^{-1}$), 100x/$O_1$-$O_3$ One-loop (-0.00534 $min^{-1}$), and 100x/$O_1$-$O_2$ One-loop (-0.00355 $min^{-1}$).

**Modeling the DNA looping**

*Description of the system.* The *lac* repressor, with two DNA binding domains, can bind two operators simultaneously by looping the intervening DNA. We consider the simplest realistic model of the *lac* operon that considers DNA looping, including the main operator $O_1$ and an auxiliary operator, referred to as $O_a$. The repressor's binding to $O_1$ prevents transcription by the RNA polymerase irrespective of its binding to $O_a$, which does not prevent transcription.

The canonical description considers that there is a set of transcriptional states $s$ and that mRNA, $m$, is produced at a rate $g_s$ for each transcription state (14). We consider explicitly 5 transcriptional states, which are labeled as follows:

| State | Description |
|---|---|
| 1 | $O_1$ and $O_a$ free |
| 2 | $O_1$ free and $O_a$ bound to the repressor |
| 3 | $O_1$ bound to the repressor and $O_a$ free |
| 4 | $O_1$ and $O_a$ bound to a repressor looping DNA |
| 5 | $O_1$ and $O_a$ each bound to a repressor |

We use a vectorial representation of the system in the state space. The transcription rates $g_s$ are expressed as the components of the vector

$$\mathbf{g} \equiv \begin{pmatrix} k_t & k_t & 0 & 0 & 0 \end{pmatrix}^T,$$

which specifies transcription taking place at a rate $k_t$ only when the main operator $O_1$ is free. Analogously, transitions between states result from the binding and unbinding of the repressor. The transition rates $k_{s,s'}$ from the state $s$ to the state $s'$ are specified through the elements of the matrix

$$\mathbf{k} \equiv \begin{pmatrix} 0 & n_R k_{\text{on}} & n_R k_{\text{on}} & 0 & 0 \\ k_{\text{off-Oa}} & 0 & 0 & k_{\text{loop}} & (n_R - 1)k_{\text{on}} \\ k_{\text{off-O1}} & 0 & 0 & k_{\text{loop}} & (n_R - 1)k_{\text{on}} \\ 0 & k_{\text{off-O1}} & k_{\text{off-Oa}} & 0 & 0 \\ 0 & k_{\text{off-O1}} & k_{\text{off-Oa}} & 0 & 0 \end{pmatrix},$$

where $k_{on}$ is the association rate of the repressor for an operator; $k_{\text{off-O1}}$ and $k_{\text{off-Oa}}$ are the dissociation rates of the repressor from $O_1$ and $O_a$, respectively; $k_{\text{loop}}$ is the rate of loop formation when the repressor is bound to one operator; and $n_R$ is the number of repressors. This description, developed originally in Ref. (15), has been shown to accurately describe the *lac* operon under an exhaustive range of experimental conditions (16, 17), with 1.7-fold accuracy over a 10,000-fold variation of the expression level.

The time evolution of the probability $P_s$ of the state $s$ is given by

$$\frac{dP_s}{dt} = \sum_{s'} [\, k_{s',s} P_{s'} - k_{s,s'}\, P_s],$$

which takes into account the transitions between transcriptional states.

The steady-state expression of the probability $P_s$ is obtained by solving $0 = \sum_{s'}[k_{s',s}P_{s'} - k_{s,s'}P_s]$, which follows straightforwardly from the preceding equation. The solution, using the statistical weights $Z_s$, is expressed in vector form as

$$\mathbf{P}^{\mathrm{ss}} = \mathbf{Z}/\|\mathbf{Z}\|_1,$$

where $\|\boldsymbol{Z}\|_1$ is the partition function expressed using the one norm.

To obtain compact expressions, we express the dissociation and the looping rates in terms of the repressor-operator association constants, $K_{\mathrm{O1}}$ and $K_{\mathrm{Oa}}$, and looping local concentration, $n_L$, as $k_{\text{off-O1}} = k_{\mathrm{on}}/K_{\mathrm{O1}}$, $k_{\text{off-A}} = k_{\mathrm{on}}/K_{\mathrm{Oa}}$, and $k_{\mathrm{loop}} = n_L k_{\mathrm{on}}$. In terms of these parameters, the statistical weights are

$$\boldsymbol{Z} \equiv (1 \quad n_R K_{\mathrm{Oa}} \quad n_R K_{\mathrm{O1}} \quad n_R n_L K_{\mathrm{Oa}} K_{\mathrm{O1}} \quad n_R(n_R-1) K_{\mathrm{Oa}} K_{\mathrm{O1}})^T.$$

The association constants and looping local concentration are related to the free energies of binding to $O_1$, $\Delta F_{\mathrm{O1}}$, and $O_a$, $\Delta F_{\mathrm{Oa}}$, and of looping, $\Delta F_L$, as $K_{\mathrm{O1}} = e^{-\Delta F_{\mathrm{O1}}}$, $K_{\mathrm{Oa}} = e^{-\Delta F_{\mathrm{Oa}}}$, and $n_L = e^{-\Delta F_{\mathrm{L}}}$, respectively, which use the thermal energy ($k_B T$) as energy units.

*Parameter values.* We use the number of molecules, abbreviated molec, as the units of substance; 1 molecule/cell, equivalent to 1.5 nM for an *E. coli* cell, as the units of concentration; and minutes, abbreviated min, as the units of time.

The association rate constant for the repressor tetramer binding to an operator at 30°C was set to $k_{\text{on}} = 0.28\ \text{molec}^{-1}\text{min}^{-1}$, consistently with the upper and lower bounds established in Fig. 2 of Ref. (7) for the repressor dimer at 37°C and 25°C, respectively.

The repressor-operator association constants and the looping concentration were obtained from the free energies inferred in Refs. (16, 17) as $K_{\text{O1}} = 2.76\ \text{molec}^{-1}$, $K_{\text{Oa}} = 0.32\ \text{molec}^{-1}$, and $n_L = 1080$ molec. These values lead to a single $O_1$ operator repression level, defined as the maximum transcription over the actual transcription and computed as $L = k_t/(\mathbf{P}^{\text{ss}} \cdot \mathbf{g})$, of $L = 27$, to a full system repression level of $L = 2291$, and to the main operator O1 being 10 times stronger than the auxiliary operator $O_2$, consistently with the experimental observations of Ref. (10), with similar growing conditions as used here but with a slightly different temperature of 32°C.

These values of $k_{\text{on}}$ and $K_{\text{O1}}$ lead to $k_{\text{off-O1}} = 0.10\ \text{min}^{-1}$, corresponding to an average occupancy time of $O_1$ by the repressor of 9.7 min, consistently with the upper and lower bounds established at 37°C and 25°C, respectively, in Fig. 2 of Ref. (7).

The transcription rate was set to $k_t = 20\ \text{molec} \cdot \text{min}^{-1}$ as reported in Ref. (18).

*Burst size and OFF interval definition.* We define the OFF interval as the time between the first binding to $O_1$ after transcription and the subsequent transcriptional event. Therefore, the OFF interval can include multiple rounds of binding and unbinding to $O_1$. Correspondingly, the ON interval is defined as the time between the first transcriptional event after unbinding from $O_1$ and the subsequent binding to $O_1$. In this way, we can unambiguously define the burst size as the number of transcripts produced during the ON interval.

To obtain analytical results, we consider a reduced description. Explicitly, we aggregate the states without potential for transcription into the bound state, denoted by B, which includes the configurations with the repressor bound to $O_1$; namely, one repressor bound to $O_1$ and the auxiliary operator free, a repressor bound simultaneously to $O_1$ and the auxiliary operator by looping the intervening DNA, and one repressor bound to $O_1$ and another one bound to the auxiliary operator. Its probability is given by $P_B = P_3 + P_4 + P_5$. Analogously, we aggregate the states with potential for transcription into the free $O_1$ state, denoted by E, which includes the configurations with the repressor not bound to $O_1$; namely, $O_1$ free and the auxiliary operator occupied, and both operators free. Its probability is given by $P_E = P_1 + P_2$.

*OFF interval statistics.* To account for transcriptional events in the reduced description, we introduce a transcription start state, denoted by TS, so that a transcript is produced when the system reaches this state. The resulting transitions between the reduced states are described by

$$B \underset{k_b}{\overset{k_u}{\rightleftarrows}} E \overset{k_t}{\rightarrow} TS,$$

where $k_b$ is the effective binding rate for the repressors to $O_1$, $k_u$ gives the unbinding rate for a repressor from $O_1$, and $k_t$ is the effective transcription rate.

When the system reaches the state E, there is a probability

$$\alpha = \frac{k_b}{k_b + k_t}$$

of returning to the bound state. The probability of $l$ unbinding events before transcription occurs is

$$P_l = \alpha^{l-1}(1 - \alpha).$$

Considering that $k_u \ll k_b$, as implied by the physical parameters of the system, the timing at which $l$ unbinding events happen is given by the composition of $l$ exponential decays, described by the Erlang distribution,

$$w_{t|l} = \frac{(k_u t)^{l-1}}{(l-1)!} k_u e^{-k_u t},$$

which results in a distribution of waiting times between transcriptional events (OFF intervals) given by

$$w_t = \sum\nolimits_l w_{t|l} P_l = \sum\nolimits_l \frac{(\alpha k_u t)^{l-1}}{(l-1)!} k_u e^{-k_u t} (1 - \alpha) = e^{-(1-\alpha) k_u t} k_u (1 - \alpha)$$

The average duration of the OFF interval is

$$\tau_{\mathrm{OFF}} = \int_0^\infty t w_t dt = \frac{1}{k_u}\left(1 + \frac{k_b}{k_t}\right)$$

More generally, the distribution of OFF intervals can be computed as the survival probability distribution of the system being in the states B and E before a transcriptional event occurs (19, 20):

$$w_t = -\frac{d}{dt}(P_B + P_E).$$

The dynamics of the corresponding probabilities is given by

$$\frac{d}{dt}P_B = -k_{on}n_L P_1 + k_{on}(n_R + n_L)P_E - k_{\text{off}-01}P_B$$

$$\frac{d}{dt}P_E = k_{on}n_L P_1 - [k_{on}(n_R + n_L) + k_t]P_E + k_{\text{off}-01}P_B$$

which forms a closed set of equations when the contribution of $P_1$ is negligible compared to that of $P_2$. In that case, we obtain the explicit values $k_u = k_{\text{off}-01}$ and $k_b = k_{on}(n_R + n_L)$. Using the initial conditions $P_B(0) = 1$ and $P_E(0) = 0$, the resulting distribution of OFF intervals is given by

$$w_t = \frac{k_u k_t e^{-\frac{1}{2}(k_u+k_b+k_t)\left(1+\sqrt{1-\frac{4k_u k_t}{(k_u+k_b+k_t)^2}}\right)t}\left(e^{(k_u+k_b+k_t)\sqrt{1-\frac{4k_u k_t}{(k_u+k_b+k_t)^2}}\,t} - 1\right)}{\sqrt{(k_u + k_b + k_t)^2 - 4k_u k_t}}$$

Since $k_u \ll k_b + k_t$, by expanding the square root, we obtain

$$w_t \simeq \frac{k_u k_t}{k_b + k_t} e^{-\frac{k_u k_t}{k_b+k_t}t}$$

which coincides with the expression obtained previously.

*Burst size statistics.* To account for multiple transcriptional events in the reduced description, we consider that, after reaching the transcription start state TS and producing a transcript, the system returns instantaneously to the free $O_1$ state E. The resulting transitions between the reduced states are described by

$$B \underset{k_b}{\leftarrow} E \underset{\infty}{\overset{k_t}{\rightleftarrows}} TS.$$

The probability of making $r$ E-TS transitions before $O_1$ is occupied is

$$P_r = (1-\alpha)^{r-1}\alpha$$

and the average number of transcripts per burst

$$\langle r\rangle = \sum\nolimits_r rP_r = \frac{1}{\alpha} = 1 + \frac{k_t}{k_b}.$$

*Stochastic simulations.* We performed stochastic simulations of the Master equation that describes the dynamics of the system under steady conditions following the approach of Ref. (14). Explicitly, the time evolution of the joint probability $P(p, m, s)$ of the number $p$ of proteins, the number $m$ of mRNA molecules, and the system state $s$ is governed by the Master equation

$$\begin{aligned}\frac{dP(p,m,s)}{dt} = &\sum\nolimits_{s'}[\,k_{s',s}P(p,m,s') - k_{s,s'}\,P(p,m,s)] + g_s[P(p,m-1,s) - P(p,m,s)] \\ &+ \lambda_m[(m+1)P(p,m+1,s) - mP(p,m,s)] \\ &+ k_p m[P(p-1,m,s) - P(p,m,s)] + \lambda_p[(p+1)P(p+1,m,s) - pP(p,m,s)],\end{aligned}$$

which takes into account the transitions between transcriptional states, mRNA production, mRNA degradation, protein production, and protein dilution. Here, $\lambda_m$ is the mRNA degradation rate, $k_p$ is the translation rate, and $\lambda_p$ is the dilution rate, which equals the growth rate. As discussed in the "Description of the system" section, $k_{s,s'}$ is the transition rate between the transcriptional states of the operon, and $g_s$ is the transcription rate in the state $s$.

The ON and OFF intervals are computed at discrete times $t_i = i\Delta t$ equally spaced by $\Delta t$ as $p(t_{i+1}) - p(t_i) > 0$ and $p(t_{i+1}) - p(t_i) \le 0$, respectively. The burst size is computed as the protein produced during an ON interval.

*Computational results.* The table below shows the average OFF interval duration, $\tau_{\mathrm{OFF}}$, and burst size, $\langle r\rangle$, computed analytically and from stochastic simulations for systems with the WT number of repressors ($n_R = 10$), with 100 times the WT number of repressors ($n_R = 1000$), with DNA looping, and without DNA looping ($K_{\mathrm{Oa}} = 0$ and $n_L = 0$):

| Looping | Repressors | $\tau_{\mathrm{OFF}}$ (min) analytical | $\tau_{\mathrm{OFF}}$ (min) simulation | $\langle r\rangle$ analytical | $\langle r\rangle$ simulation |
|---|---|---|---|---|---|

| | | | | | |
|---|---|---|---|---|---|
| Yes | WT | 162 | 160 (154) | 1.1 | 1.5 (1.4) |
| Yes | 100×WT | 299 | 310 (305) | 1.0 | 1.1 (1.1) |
| No | WT | 11.1 | 22.5 (13.3) | 7.9 | 23.3 (13.0) |
| No | 100×WT | 149 | 159 (153) | 1.1 | 1.2 (1.1) |

The parameter values for the analytical computations are described in the "Parameter values" section. Additional parameters in the simulations include $\lambda_m = 0.33\ \text{min}^{-1}$, $\lambda_p = 0.0059\ \text{min}^{-1}$, and $k_p = 26.7\ \text{min}^{-1}$. Averages from simulations were computed over $1.9\times10^6$ min runs after discarding the initial $3.9\times10^5$ min. Simulation results were computed by sampling the time series every 5 min (results without parentheses) or 1 min (results within parentheses). The average burst size from the stochastic simulations in the previous table has been normalized by the average protein produced by a single transcript.

**Simulation with cell division**

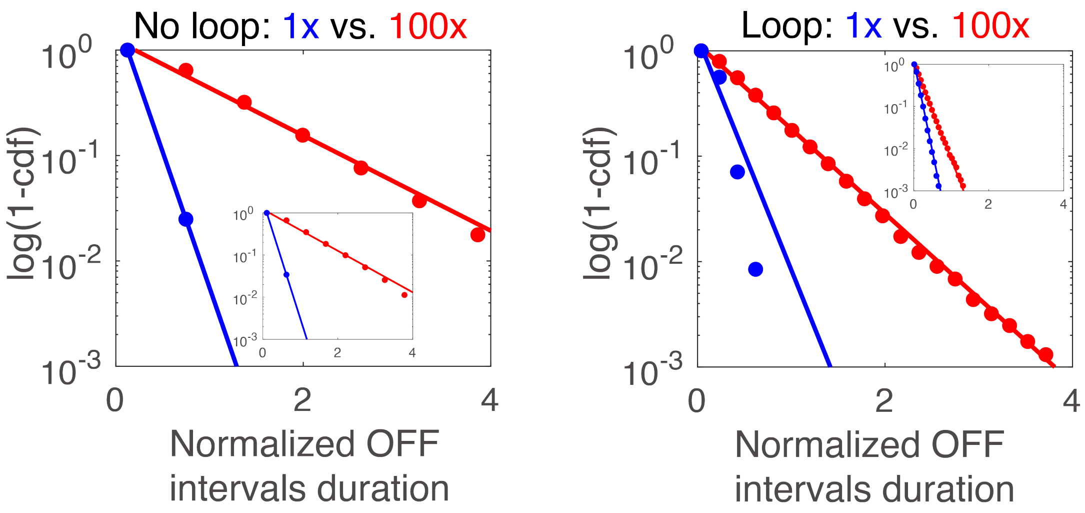

**Fig. S14. Statistics of OFF intervals in simulation with cell division.** We calculated the cumulative distributions of OFF intervals for simulations with cell division by forcing the repressor to unbind from the operators periodically. The insets give the simulations without cell division, which are identical to that in Fig. 2c. Normalization factors: Loops strains with cell division (515 min), Loops strains without cell division (1685 min), No-loop strains with cell division (160 min), and No-loop strains without cell division (190 min).

To assess the effects of cell division on the repression of the *lac* promoter, we performed stochastic simulations that differ from those in the insets of Fig. 2c by forced periodical unbinding events of the repressor from the operators (Fig. S14). For the Loops strain, the statistics of the OFF intervals in the presence of cell division deviates from that of an exponential distribution. The mean interval duration is half shorter in the presence than in the absence of division. Furthermore, the OFF intervals in the presence of cell division are sensitive to repressor concentration changes and shows a four-fold reduction. On the other hand, the effect of cell division on the No-loop strains is very mild.
